# Chronic alteration of Ca^2+^ and hemodynamic signals induced by intracortical microstimulation in the visual cortex of awake mice

**DOI:** 10.1101/2025.10.23.679242

**Authors:** Naofumi Suematsu, Alberto L Vazquez, Takashi DY Kozai

## Abstract

Understanding the chronic effects of intracortical microstimulation (ICMS) and device implantation on cortical function is essential for the development of stable neuroprosthetics. We chronically implanted microelectrode into Thy1-GCaMP6s mice and conducted longitudinal mesoscopic widefield and two-photon Ca^2#x002B;^ imaging alongside intrinsic optical signal recordings over 12 weeks. Six ICMS frequencies (10-500 Hz) and contralateral visual LED stimuli were delivered in repeated sessions. Oxygen extraction fraction (OEF) was estimated from dual-wavelength reflectance, and hemodynamic response functions (HRF) were derived via regularized deconvolution. Over the chronic period, 25-Hz ICMS evoked progressively larger Ca^2+^ responses whose duration was extending, while spatial spread remained stable around 500 µm. Concurrently, OEF reductions deepened, reflecting increased relative blood supply, and the spatial extent of OEF signals contracted in the first week before expanding by day 21. After discharge-like Ca^2+^ events emerged in nearly half of mice, predominantly under 25-Hz ICMS. HRF peak amplitude increased between day 0 and days 7-21, with latency decreasing. Chronic ICMS induces progressive potentiation of neuronal and vascular responses alongside local neurovascular uncoupling on day 0 and sustained silencing of spontaneous activity near the implant. These dynamics exhibiting a stabilization 6-7 weeks after the implantation, coupled with frequency-dependent after discharges, underscore the need for adaptive stimulation strategies prior to the stabilization and to prevent the after discharge and implant designs that mitigate local suppression and ensure long-term stability of cortical neuroprostheses.

## 1.0 Introduction

Penetrating microelectrodes have long been instrumental in advancing neurophysiological research, enabling precise recording^1-4^ and stimulation^5-8^ of neural activity. These technologies have shown significant promise in neuroengineering applications, particularly in the development of neuroprosthetics^9-11^. Intracortical microstimulation (ICMS), delivered through implanted probes in the cortex, has been extensively used to evoke neural responses^12-14^, influence behavior^15-18^, and even restore aspects of sensory perception^19-23^, such as vision, in human subjects. Despite these advances, ICMS efficacy often degrades over time due to chronic tissue responses surrounding the implant, including gliosis, fibrotic encapsulation, and vascular remodeling^1,24^.

The long-term success of any implanted neural interface depends on its functional biocompatibility, which is defined as the ability of the device to perform its intended function, with the desired degree of integration in the host, without eliciting any undesirable local or systemic effects in that host^25^. For recording interfaces, biocompatibility has been extensively studied, focusing on how the foreign body response impacts chronic signal quality^1,26-34^. In contrast, the biocompatibility of ICMS devices, their ability to reliably drive neuronal networks over time despite ongoing tissue responses, remains poorly characterized. While acute optical tools in combination with ICMS have characterized neuronal responses within hours of implantation^5,35-49^, these data do not capture how stimulation efficacy progresses as the interface transitions from the acute to chronic phase.

Acute trauma from implantation causes vessel rupture^50^, mechanical strain of neuronal and glial bodies^31,51-53^, protein adsorption^32^, yet early ICMS responses often characterized by robust neuronal activation^40^. Most cellular-level ICMS studies have focused on this acute period^35,39-44,46-48,52^. Over time, however, these acute disruptions^50,54-57^ advance into a chronic foreign body response that can progressively impair stimulation efficacy. Key inflammatory and metabolic stress processes include the formation of foreign body giant cells^58^, fibrotic encapsulation^59-64^, lysosome dysfunction^56^, neutrophil infiltration^1,58^, monocyte differentiation into macrophages^1,65^, vascular remodeling^38,54,66^, and ischemia^24^, all of which disrupt local neural activity and degrade the electrode-tissue interface. Fibrotic encapsulation reduces current spread and elevates stimulation threshold over time^67^. Concurrently, local vasculature remodeling^38,54^, associated with altered blood flow and tissue oxygenation^1^, may exacerbate inflammation related elevated metabolic demand^56,57^, leading to long-term neuronal depression^40^ and altered neuronal excitability^24,68,69^.

The relationship between ischemia and ICMS-driven activity is complex. While local neurons in ischemic zones may be hypoactive^1^, ICMS can still activate passing axons and elicit antidromic propagation to distant, metabolically healthy somas. Consequently, ICMS-evoked activity in chronically implanted cortex may not accurately reflect local tissue viability, complicating interpretation and control of stimulation-evoked percepts.

Moreover, neuronal and network excitability progresses dynamically as the implantation injury resolves, and ICMS itself may further modulate excitability or increase metabolic^69-73^. Characterizing this post-implantation transition, from acute injury to chronic adaptation, is essential for assessing long-term stimulation efficacy. Understanding ICMS biocompatibility therefore requires longitudinal, multimodal visualization of both neuronal and vascular responses under repeated stimulation, an area where data remain limited^74^.

Recent advances in optical imaging are beginning to clarify these chronic dynamics by providing detailed spatial and temporal insights into neural activity and hemodynamic responses^40^. Widefield fluorescence imaging and two-photon microscopy allow longitudinal monitoring of Ca^2+^ dynamics and blood flow in the cortex^75^, offering a deeper understanding of how chronic tissue responses impact neuronal function.

Complementary intrinsic signal imaging can further reveal cortical oxygenation changes and metabolic stress. Together, these techniques offer high spatiotemporal resolution to determine whether chronic ICMS degradation arises from mismatches between metabolic supply and stimulation-driven demand.

Here, we investigate how chronic tissue responses to microelectrode implantation alter spontaneous, sensory-evoked, and ICMS-evoked cortical activity in the primary visual cortex over time. Using Thy1-GCaMP6s mice, which express a genetically encoded Ca^2+^ indicator in excitatory neurons, we performed *in vivo* longitudinal imaging over 12 weeks post-implantation to track evolving patterns of neuronal and hemodynamic responses. We hypothesize that chronic tissue responses will progressively diminish ICMS efficacy by altering the magnitude and spatial pattern of neuronal activation and suppression. By directly visualizing these changes, this study fills a critical gap in understanding how the foreign body response shapes the functional lifetime of stimulation-based neural interfaces.

## 2.0 Methods

All procedures followed our previous study^40^, here we briefly describe these procedures.

### 2.1. Animals and surgery

Transgenic mice expressing the Ca^2+^ indicator in excitatory neurons (Thy1-GCaMP6s; 024275, Jackson Laboratory, ME, USA) were used (n = 9, same mice were used both for widefield imaging and two-photon imaging, see Table S1 for details). Surgical procedures were conducted following previously established protocols^42-44^. Mice were anesthetized with a mixture of xylazine (7 mg/kg body weight [b.w.], intraperitoneally [i.p.], Covetrus, OH, US) and ketamine hydrochloride (75 mg/kg b.w., i.p., Covetrus). Craniotomies were performed over the left visual cortex (AP = -1 to -5, LM = 0.5 to 3.5 mm), which were larger than the imaging area. Custom stimulation microelectrodes (Q1×4-3mm-50-703-CM16LP, NeuroNexus Technologies, single 3-mm-long shank, 30-µm-diameter four channels, 50-µm site spacing, electrochemically activated iridium/iridium oxide sites^76^) were implanted at an angle of 20-30 degrees slanted towards the posterior region. The electrode was secured to the skull with UV-curable resin (e-on Flowable A2, BencoDental, PA, USA). Reference and ground of the electrode were wired to the bone screws placed in the skull over the frontal lobes. The exposed cortical surface was covered with Kwik-Sil and a cover glass, which was glued with the UV-curable resin.

Anesthetic depth was periodically assessed by pedal withdrawal reflex and respiratory rate. Respiration was monitored visually by chest movement. Supplemental ketamine (45 mg/kg b.w., i.p.) was administered as needed to maintain anesthetic depth. Following surgery, atipamezole hydrochloride (1 mg/kg b.w., i.p., Antisedan, Zoetis, NJ, US) and ketoprofen (5 mg/kg b.w., i.p., Ketofen, Zoetis) were administered for reversal of anesthesia and for analgesia. The mouse body temperature was maintained using a heating water pad and respiration was monitored visually from the beginning of the surgery until recovery (< 2hr). In the following two days after surgery, additional ketoprofen was administered daily. Animals were kept in a 12-h/12-h light/dark cycle. All procedures described here were approved by the Division of Laboratory Animal Resources and Institutional Animal Care and Use Committee at the University of Pittsburgh and performed in accordance with the National Institutes of Health Guide for Care and Use of Laboratory Animals.

### 2.2. Mesoscopic and Microscopic Image Acquisition

Chronic recordings were conducted over multiple days after surgery (Fig. 1A). Each recording day, animals were placed in a custom treadmill and head-fixed throughout the imaging experiments to minimize motion artefacts while allowing voluntary locomotion. We employed a mesoscopic-scale widefield microscope (MVX-10 epifluorescence microscope, Olympus, Tokyo Japan) for simultaneous recording of Ca^2+^ fluorescent and intrinsic optical signals^40,75,77^. The recording camera (QImaging Retiga R1, Cairn Research Ltd, Kent, UK) was controlled using Micro-Manager software^78,79^. Illumination and acquisition of blue (for GCaMP excitation, 482.5±15.5 nm, #67-028, Edmund Optics Ins., NJ, US), green (for total blood volume, 532±2 nm, FLH532-4, Thorlabs Inc., NJ, US), and red (for differentiation of oxy- and deoxy-hemoglobin concentrations, 633±2.5 nm, FLH633-5, Thorlabs Inc.) channels were controlled sequentially. Fluorescent (GCaMP) and reflectance images (hemoglobin) were long-pass filtered (> 500 nm, ET500lp, Chroma Technology Corporation, VT, US) before acquisition.

**Figure 1.**
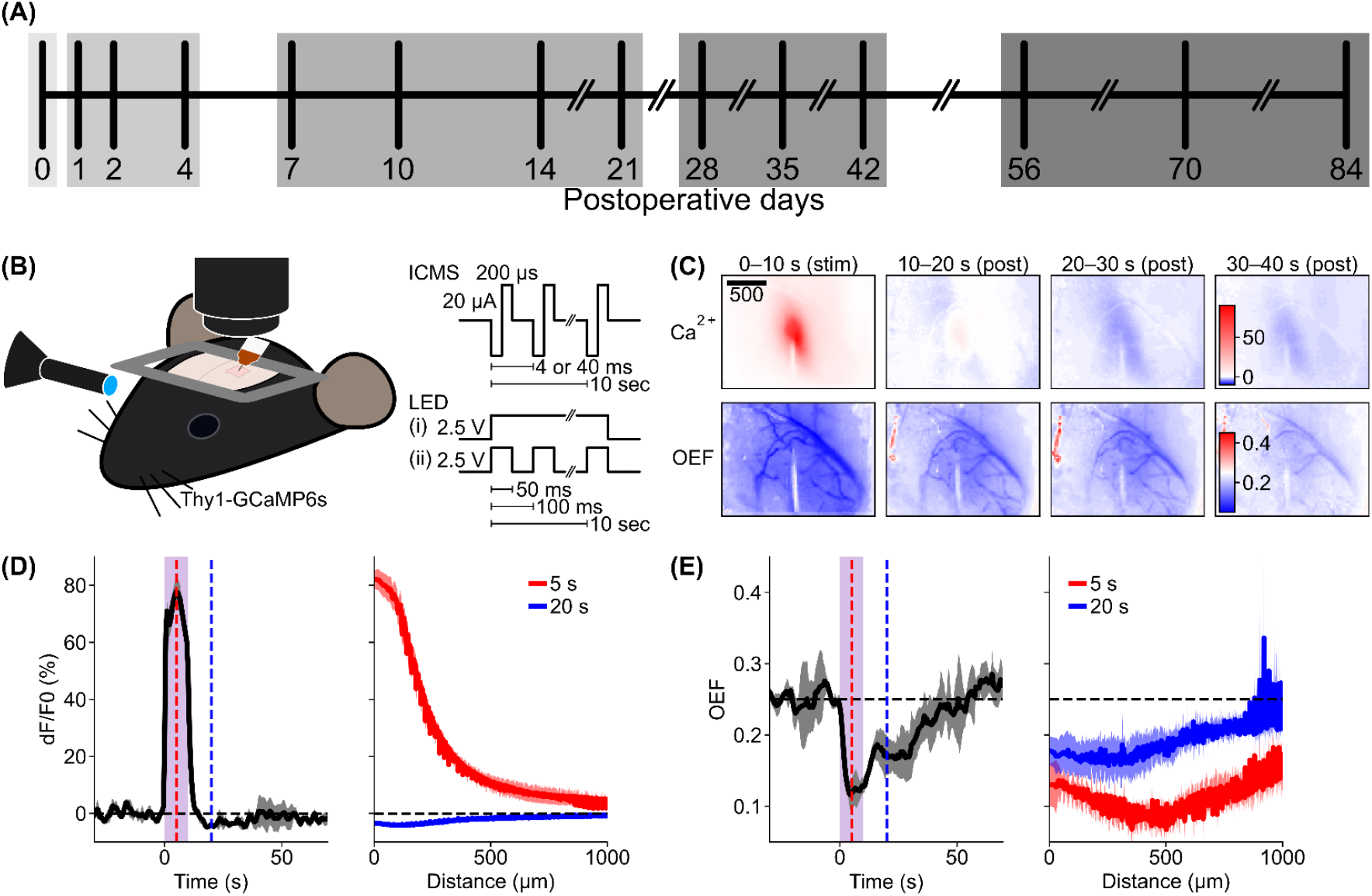
Chronic assessment of visual cortical responses to ICMS and visual stimulation. (**A**) Timeline of chronic recordings. (**B**) Imaging and stimulation setups. Widefield and two-photon imaging was performed on the implanted hemisphere, and visual stimulation was applied to the contralateral eye. (**C**) Example widefield images of neural (Ca^2+^ dF/F_0_) and OEF hemodynamic responses induced by 25-Hz ICMS. Scale bar = 500 µm. (**D and E**) Time course (left) and distance profile (right; measured at 5 [red] and 20 [blue] seconds after stimulation onset) of induced neural (D) and hemodynamic (E) responses. Mean ± SD across four trials. Shaded bar indicates stimulation period (0-10 s); vertical dashed lines mark the time points of distance profiles.

Two-photon microscopy (Bruker, Madison, WI, US) was performed using a laser tuned to 920-nm wavelength (Insight DS+, Spectra-Physics, Menlo Park, CA, US) for GCaMP6s excitation (laser power ≈ 10-20 mW). Resonant Galvo scanning was performed to capture higher resolution images at 30 Hz over a 413 × 413 µm^2^ field of view (512 × 512 pixels) at a single z-plane focused around the stimulation site (150-200 µm below pia). In both imaging sessions (mesoscopic-scale and two-photon imaging), each recording was composed of a 30-s pre-stimulation period, a 10-s stimulation period, and a 60-s post-stimulation period, which was repeated 28 times (see below). Both imaging sessions were operated for each recording day (see Table S1 for details).

### 2.3. Electrical (ICMS) and Visual Stimulation

ICMS was performed using an IZ2 stimulator controlled by an RZ5D base processor (Tucker-Davis Technologies, FL, US). Each pulse consisted of a 200-µs cathodic leading phase followed by a 200-µs anodic phase, with balanced charges between the cathodic and anodic phases. The current was fixed to ±20 µA (4 nC/phase), selected to be lower than the safety limit^80,81^ (electrode impedance ≈ 100–200 kΩ at 1 kHz). Six different train frequencies (10, 25, 50, 100, 250, and 500 Hz) were tested. The order of ICMS trials was repeated two times in ascending-descending frequency order (10 to 500, 500 to 10, 10 to 500, 500 to 10 Hz).

For comparison, a single blue LED was positioned in front of the animal’s eye contralateral to the imaging window as a more natural visual stimulus (n = 5-9 mice depending on days, see Table S1 for details). The LED was controlled by the stimulator, applying a constant current or square wave at 10 Hz (50-ms on, 50-ms off, 50% duty cycle) with an amplitude of 2.5 V. All stimulation trials were repeated 4 times. Both ICMS and visual stimulation continued for 10 s.

### 2.4. Analyses and Statistics

To facilitate analysis across biologically meaningful timepoints, recording sessions were divided into five groups based on days after surgery (Fig. 1A): day 0, days 1-4, days 7-21, days 28-42, and days 56-84. These timepoints were selected to align with distinct phases of the neural tissue response following microelectrode implantation^60,82^. Day 0 represents the immediate effects of surgical insertion, including mechanical disruption, ischemia, and the onset of acute neuronal and glial stress. Days 1-4 capture the acute injury response phase, characterized by blood-brain barrier disruption, neutrophil infiltration, and the initiation of microglial activation. Days 7-21 correspond to the peak inflammatory and fibrotic induction phase, marked by maximal glial reactivity, macrophage infiltration, and the onset of extracellular matrix deposition. Days 28-42 reflect an intermediate stage during which the inflammatory response begins to transition into a chronic immune response, but fibrotic encapsulation and vascular remodeling continue to progress. Finally, days 56-84 represent the chronic stable phase, during which tissue responses plateau and long-term effects on neuronal excitability and stimulation efficacy become more apparent. These temporal groupings align with known histopathological phases of the electrode-tissue response, facilitating interpretation of functional changes across implantation time^31^.

#### 2.4.1. Pre-processing of the widefield and two-photon imaging data

Analyses were performed using custom-made scripts written in Python (ver. 3.9, Python Software Foundation, DE, US). Spatial (20 µm/pix for mesoscopic-scale, 1 µm/pix for two-photon) and temporal (2 Hz) binning with Gaussian anti-alias filtering were applied to improve signa-to-noise ratio. Image registration was conducted to align frames to the first frame of each acquisition and/or across conditions by cross-correlation with sub-pixel motion estimation^83^. For the two-photon microscopy images, additional inter-line alignment was applied as necessary to correct for shearing between lines.

For the widefield imaging data, masks were generated to exclude blood vessels as proposed previously^40^. This procedure also removed probe shank pixels that exhibited noticeably low fluorescence and reflectance values.

#### 2.4.2. Somatic region-of-interest (ROI) definition for the two-photon imaging data

Response images of excitatory neurons acquired under the two-photon microscopy were constructed by calculating pixel-wise mean fluorescent change ratio during the stimulation period (0-10 s) over pre-stimulation mean fluorescent intensity across the entire imaging region. For each animal and day, these response images for all stimulation conditions were averaged. Region of interest (ROI) for each soma was manually defined with Fiji^84^ (ImageJ 1.54d). All other pixels outside the somatic ROIs were defined as a neuropil ROI.

#### 2.4.3. Neuronal Ca^2+^ response quantification for widefield and two-photon imaging

To quantify the excitatory Ca^2+^ activity for two-photon imaging, the fluorescence change ratio (dF/F_0_) was calculated as a pixel-wise manner:

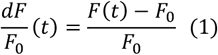

where F(t) represents the fluorescent intensity at time t and F_0_ is the mean fluorescent intensity over the pre-stimulation period (-30 s ≤ t < 0 s). For analysis relative to the electrode distance, pixels were later binned by distance from the electrode to generate distance-dependent profiles or time courses.

In the case of the mesoscopic-scale widefield imaging, the Ca^2+^ signals were corrected based on the green channel values to account for the hemodynamic effect that could potentially affect the green GCaMP signals ^75,77^. The corrected dF/F_0_(t) was calculated using the formula:

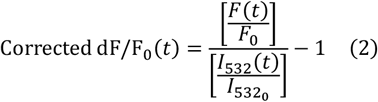

where *I*_532_(t) represents the reflectance intensity to the green illumination at a specific time point *t*, and 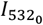 is the mean of the reflectance intensity to the green illumination during the pre-stimulation period. Distance information (defined radially from stimulation site) was subsequently incorporated to generate distance-dependent activity profiles, allowing assessment of spatial propagation of Ca^2+^ responses relative to the electrode.

#### 2.4.4. Hemodynamic activity quantification of mesoscopic-scale widefield imaging

The oxygen extraction fraction (OEF; relative concentration of deoxyhemoglobin in total hemoglobin concentration) was estimated based on the pixel-wise reflectance values in the green and red channels under the mesoscopic-scale imaging using the following equation with several assumptions of steady-state hemoglobin concentrations^40,75^:

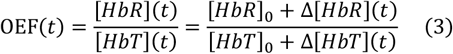

where Δ[HbT](t) and Δ[HbR](t) indicate the concentrations of total hemoglobin and deoxyhemoglobin at a time point t, respectively. [HbR]_0_ and [HbT]_0_ represent assumed baseline concentrations of the deoxyhemoglobin (15 µM) and total hemoglobin (60 µM). By definition, OEF values range between 0 and 1, and estimated OEF values outside this range (1.6 ± 1.1% pixels, mean ± standard deviation [SD] across all animals) were clipped to the valid range [0, 1] to maintain physiological interpretability by preserving polarity of response (increase vs. decrease).

To define the activation and depression distances (from stimulation site), latencies (onset timing), and durations (offset timing - onset timing) of the spatiotemporal magnitude profile of the corrected dF/F_0_ and OEF while reducing the effect of data fluctuations, we employed the hysteresis thresholding as described previously^40^.

After defining the distance, latency, and duration of significant Ca^2+^ activation or depression and hemodynamic activities, these measurements were averaged across time for each mouse. Also, the averaged measurements for each mouse were averaged across mice to evaluate these effects as a function of stimulation conditions such as ICMS frequencies and visual stimulation.

#### 2.4.5. Clustering spatiotemporal dynamics of mesoscopic-scale Ca^2+^ responses

Each activation data was clustered into several groups by a hierarchical clustering method (“AgglomerativeClustering” implemented in scikit-learn, Python module) based on its magnitude, distance, and duration, to identify episodes consistent with epileptiform-like activity characterized by abnormally strong activation. When clustering, the number of clusters was not predefined, and distance between clusters was assumed at least 10.

For visualization of the clustering results by summarizing the induced activation properties with the criteria of epileptiform activity, uniform manifold approximation and projection (UMAP; implemented in umap, Python module) was applied to the dataset of activation magnitude, distance, and duration, which were mapped into a two-dimensional space.

### 2.4.6. Hemodynamic response function

Oxyhemoglobin concentration change (dHbO; vector **O**; Fig. S13) is assumed to be represented as a convolution of neural activity (i.e. Ca^2+^; matrix **C**) and hemodynamic response function (HRF; vector **H**):

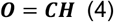

Instead of a direct deconvolution for estimating the HRF, we calculated a diagonal loading least-square deconvolution^75^ to minimize the effects of noisy fluctuations contaminated in the signals:

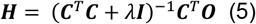

where ***I*** indicates an identity matrix. The loading weight λ was introduced as a regularization parameter and set to 0.1 (empirically chosen) throughout our study. Signals were five-times upsampled to 10 Hz (originally 2 Hz) with polyphase filtering before calculating the deconvolution. Pre-stimulation signals were trimmed to quantify the stimulation-induced hemodynamic coupling. When estimating the hemodynamic response functions from Ca^2+^ and OEF signals instead of dHbO (Fig. S14), the baseline value of OEF (assumed to be 0.25) was subtracted, followed by inverting the sign (increase in oxygen consumption from pre-stimulation baseline corresponds to increase in the pre-processed OEF signals), that helps easier interpretation of the reconstructed hemodynamic response functions (referred as OERFs). This subtraction-inversion process ensured that positive deflections correspond to increased oxygen consumption.

#### 2.4.7. Statistical procedures

For one-sample comparison with a specific value, one-sample t-test (alternative hypothesis: sample mean is different from a specific value [two-tailed]) was used with Bonferroni correction (Fig. 6B and D, number of comparisons = 5 day-periods at each distance bin). When evaluating chronic transitions of parameters for each stimulation condition, we used a linear mixed-effects (LME) model (fixed-effect variable = day, intercept and slope were varied by each subject [e.g. mouse]), followed by two-tailed paired t-test for mouse data (e.g. Fig. 2D) or two-tailed Welch’s t-test for cellular data (e.g. Fig. 4B) as post-hoc tests to compare with days 0 or 84 with Holm-Bonferroni correction (number of comparisons = 13 [14 recording days - 1]). One-way analysis of variance (ANOVA) was used for a comparison among three (or more) groups, followed by post-hoc Tukey’s honestly significant difference (HSD) test if significant difference was observed by ANOVA (e.g., Fig. S1A). Statistical significance was evaluated based on a significance level of *p* < 0.05. Significant results are denoted in the figures and summarized in supplementary tables (Table S2 for main figures, Table S3 for supplementary figures).

**Figure 2.**
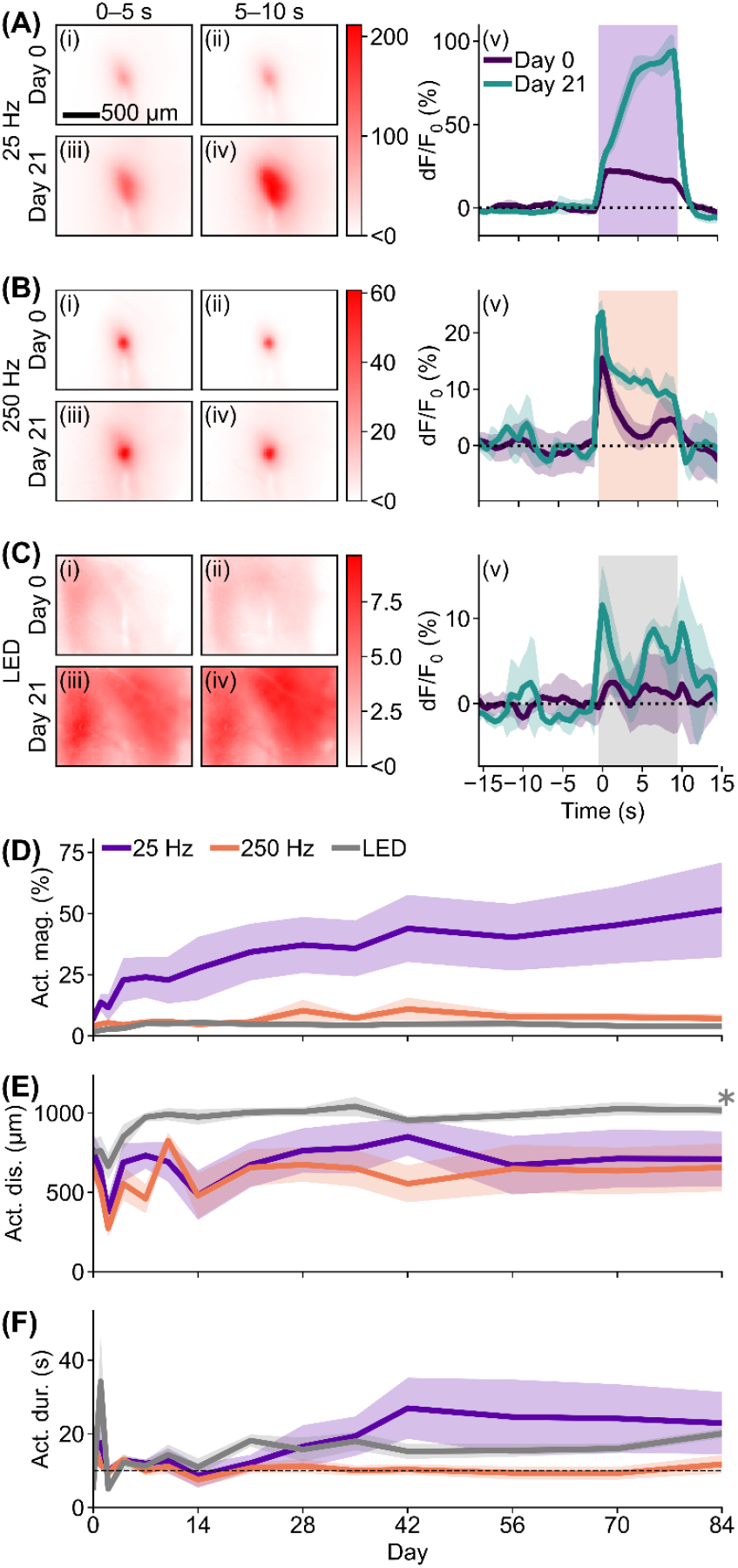
Ca^2+^ activation progressively increased after probe insertion. (**A-C**) Example colormaps from a single mouse showing Ca^2+^ responses during (**A**) 25-Hz ICMS, (**B**) 250-Hz ICMS, and (**C**) visual stimulation at (i and ii) day 0 and (iii and iv) day 21. Only increases from pre-stimulation baseline are shown in red. Corresponding time courses within 500 µm from the stimulation site are shown in (v) (mean ± SD across 4 trials). (**D-F**) Chronic comparisons of Ca^2+^ activation magnitude (D), spatial spread (E), and duration (F) across days post-insertion. Mean ± SEM across mice. ^*^ next to line plot indicates a significant effect of day (LME model).

**Figure 3.**
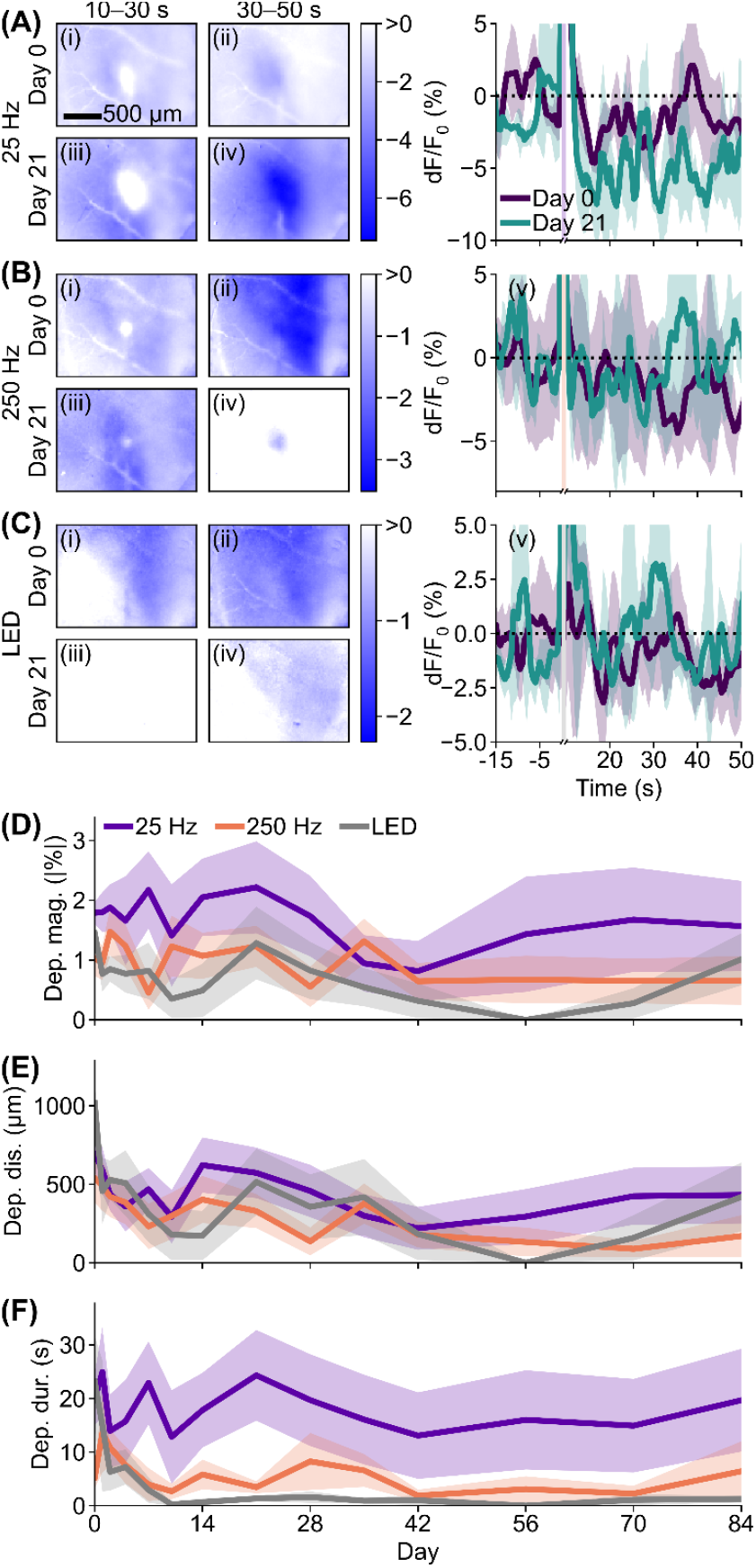
Chronic changes in Ca^2+^ depression following probe insertion. (**A-C**) Example colormaps of a single mouse showing Ca^2+^ responses after (**A**) 25-Hz ICMS, (**B**) 250-Hz ICMS, and (**C**) visual stimulation at (i and ii) day 0 and (iii and iv) day 21. Only decreases from pre-stimulation baseline are shown in blue; darker blue indicates stronger depression. Corresponding time courses within 500 µm from the stimulation site are shown in (v) (mean ± SD across 4 trials). Traces during the stimulation period (0-10 s) are omitted for clarity. (**D-F**) Chronic comparisons of Ca^2+^ depression magnitude (D), spatial spread (E), and duration (F) across days post-insertion. Mean ± SEM across mice.

**Figure 4.**
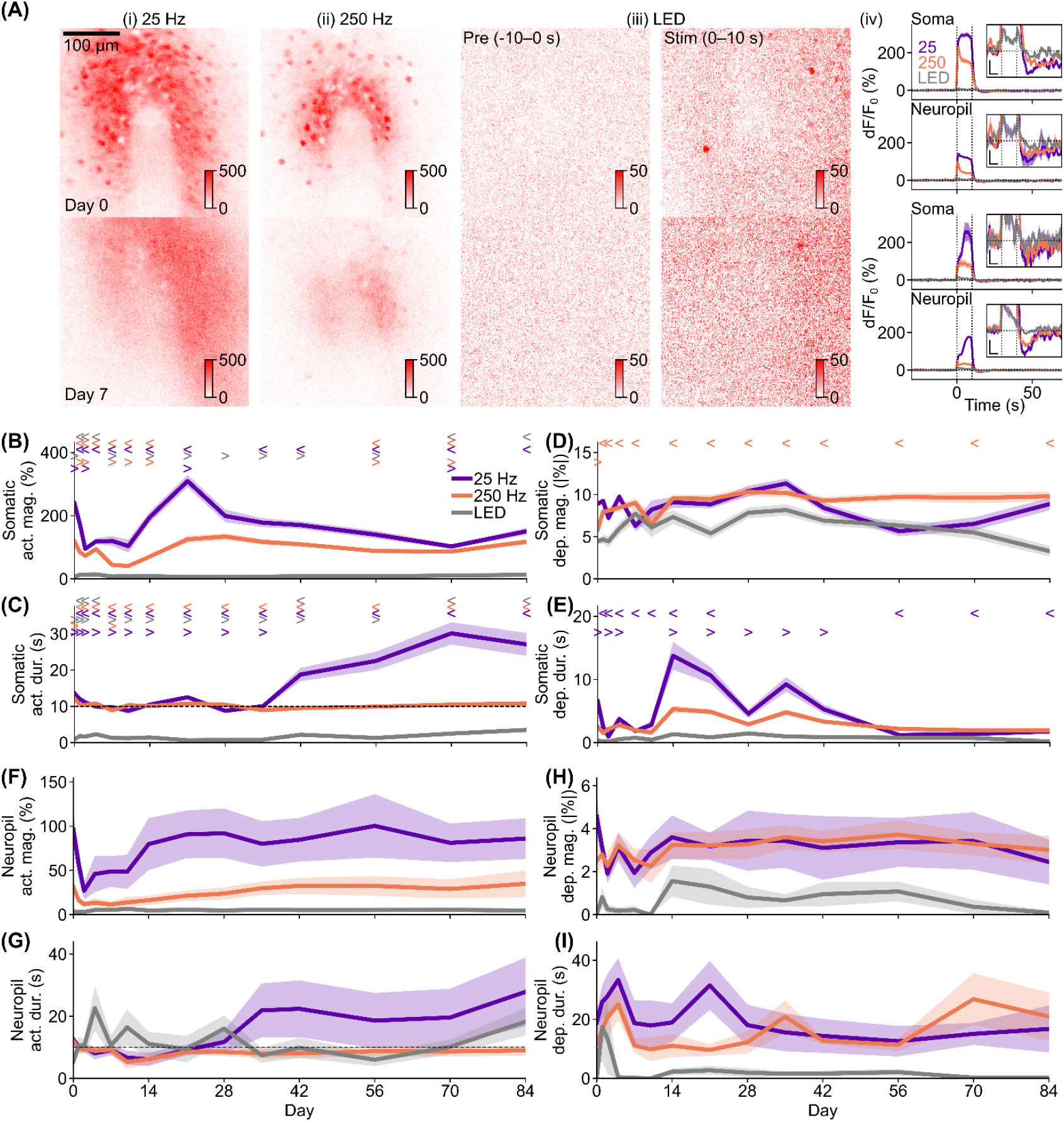
Two-photon imaging revealed chronic activation impairment-recovery trends in soma and neuropil and depression fluctuation depending on stimulation condition in soma. (**A**) Example Ca^2+^ response images at days 0 and 7 under (i) 25-Hz ICMS, (ii) 250-Hz ICMS, and (iii) visual stimulation. Corresponding time courses are shown in (iv; mean ± SEM across cells for soma, mean ± SEM across repeats for neuropil; scale bars = 5 s and 5%). (**B and C**) Chronic comparisons of somatic activation magnitudes (B) and durations (C). (**D and E**) Somatic depression magnitude (D) and duration (E) across days. (**F and G**) Neuropil activation magnitude (F) and duration (G) across days. (**H and I**) Neuropil depression magnitude (H) and duration (I) across days. Mean ± SEM across cells (B-E) or mice (F-I); “<“ and “>“ indicate significant differences compared with days 0 and 84, respectively (LME model followed by Welch’s t-test with Holm-Bonferroni correction).

**Figure 5.**
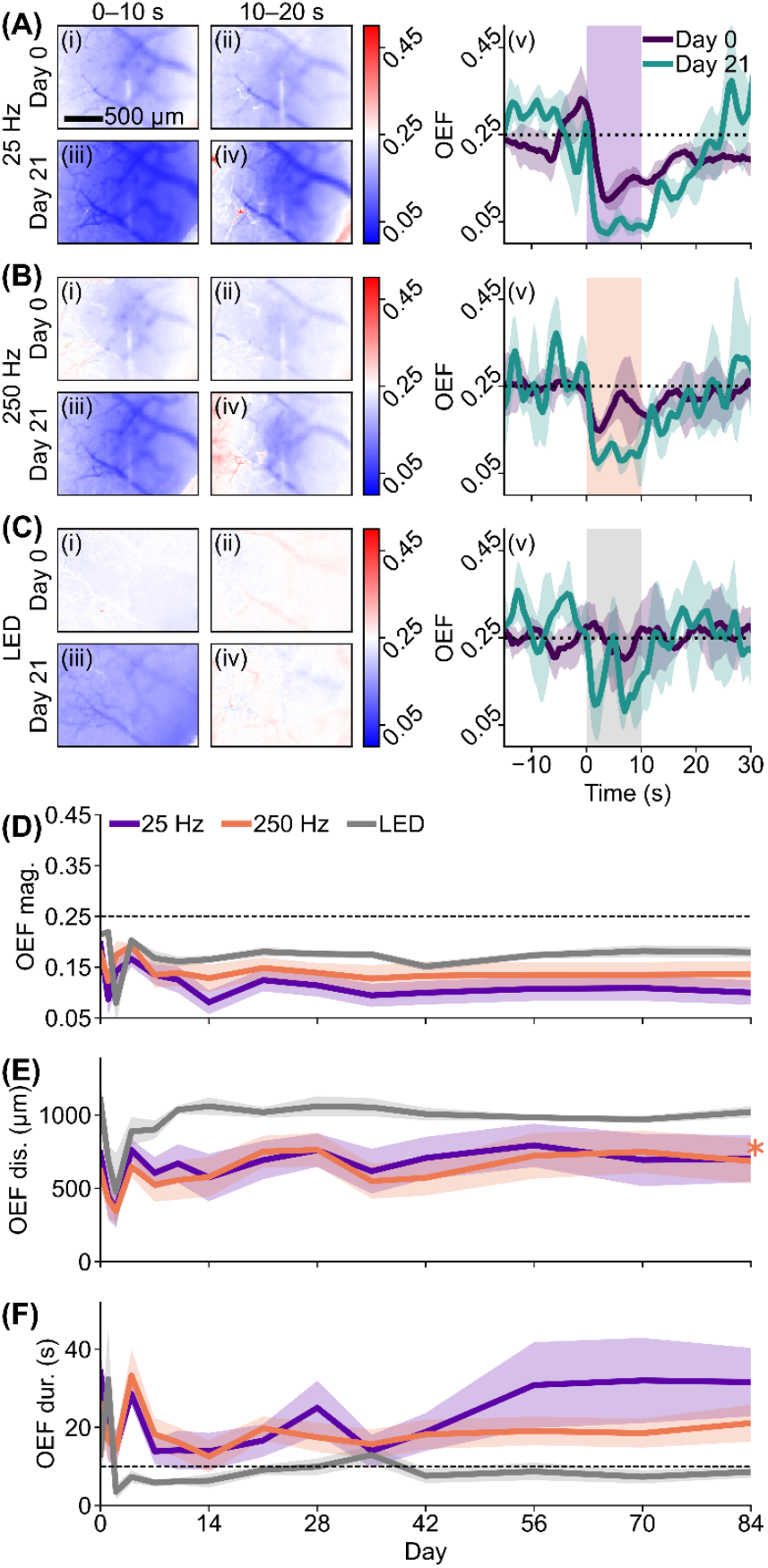
Chronic OEF decline implies metabolic supply recovery after probe insertion. (**A-C**) Example OEF response colormaps from a single mouse during and after (**A**) 25-Hz ICMS, (**B**) 250-Hz ICMS, and (**C**) visual stimulation at (i and ii) day 0 and (iii and iv) day 21. Corresponding time courses within 500 µm of the stimulation site are shown in (v; mean ± SD across 4 trials). OEF represents deoxyhemoglobin relative to total hemoglobin (see 2.4.4), where decreases from baseline (0.25) indicate relative increases in oxyhemoglobin, reflecting enhanced metabolic supply. (**D-F**) Chronic comparisons of OEF magnitude (D), spatial spread (E), and duration (F) across days post-implantation. Mean ± SEM across mice. ^*^ next to line plot indicates a significant effect of day (LME model).

**Figure 6.**
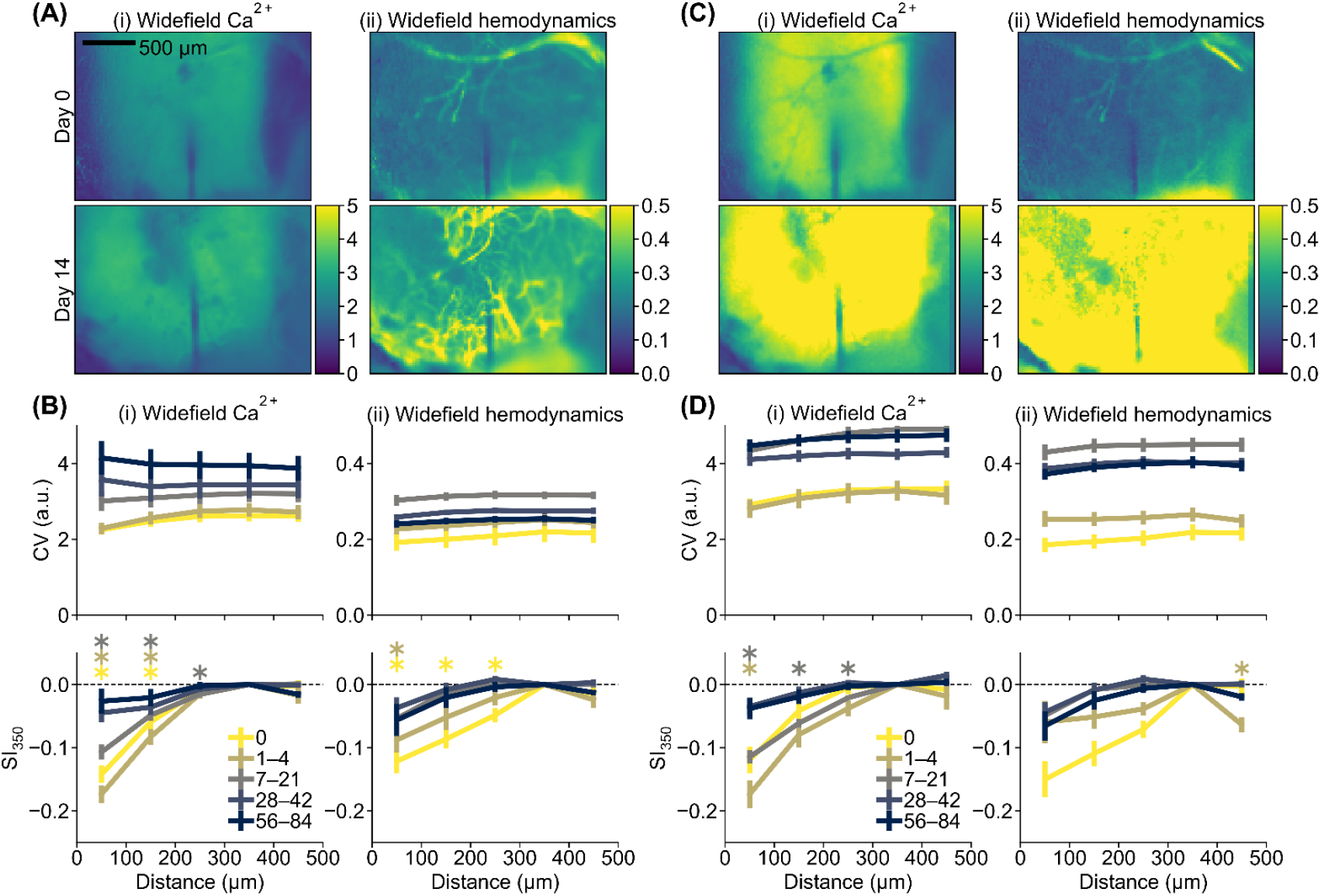
Chronic probe-induced silencing of spontaneous neural and hemodynamic activity. (**A**) Coefficient of variation (CV, pixel-wise standard deviation normalized by mean over time) of widefield Ca^2+^ (i) and hemodynamics (ii) during pre-stimulation periods at day 0 (top) and day 14 (bottom). (**B**) CV (top) and silencing index relative to 350 µm (SI_350_ = [CV_d_-CV_350_]/CV_350_, where d = 50, 150, 250, 350, or 450 μm) during pre-stimulation periods (bottom) across distances and imaging days. (**C**) CV map during visual stimulation. (**D**) CV and SI_350_ during visual stimulation. ^*^ in SI_350_ plots indicate significant differences from 0 (one-sample t-test with Bonferroni correction, *p* < 0.05; colors correspond to days after implantation).

## 3.0 Results

To investigate how ICMS modulates both neural and vascular activity over chronic timescales, we performed longitudinal widefield optical and two-photon calcium imaging in awake Thy1-GCaMP6s mice implanted with custom four-channel microelectrode arrays in primary visual cortex. Across a 12-wk post-implantation period, we delivered repetitive ICMS pulse trains spanning a physiologically relevant frequency range (10-500 Hz) and recorded mesoscale network and single-cell activity, alongside intrinsic optical signals used to quantify concurrent hemodynamic responses (**Fig. 1**). Widefield calcium signals were corrected for wavelength-dependent vascular artifacts, and regional oxygen extraction fraction (OEF) was computed from simultaneous green and red reflectance changes. Hemodynamic response functions (HRFs) were then estimated using regularized temporal deconvolution of deoxygenated hemoglobin (dHbO) signals with local calcium activity to assess neurovascular coupling. We extracted spatial and temporal features of both activation and suppression, quantified their evolution across acute, subacute, and chronic stages, and directly compared ICMS-evoked responses with sensory, visually driven activity patterns. Together, this integrated framework enabled longitudinal mapping of excitatory neuronal and vascular co-dynamics during chronic ICMS, providing quantitative insight into the functional biocompatibility of implanted stimulation devices, defined here as the sustained capacity to evoke reliable network activation without inducing maladaptive tissue responses over time.

### 3.1 Stimulation-induced Ca^2+^ activation chronologically increased after probe insertion

First, we investigated how ICMS-evoked Ca^2+^ activation progress over days following probe insertion (Fig. 2) as an indicator of functional biocompatibility. In an example animal, both 25-Hz and 250-Hz ICMS (Fig. 2A and B, respectively) and visually evoked stimulation (Fig. 2C) produced stronger cortical Ca^2+^ activation on day 21 compared to day 0. Day 0 served as the acute baseline prior to stabilization of inflammatory and foreign body responses. These two ICMS frequencies represent distinct operational regimes: 25-Hz ICMS corresponds to the frequency eliciting maximal somatic and neuropil activation in our previous work^40^, and 250-Hz ICMS approximates the upper bound commonly used in clinical applications^11,85^.

Longitudinal quantification showed a progressive increase in Ca^2+^ activation magnitude after probe insertion (Fig. 2D; Fig. S1Ai, Bi, and Ci). For the spatial spread, ICMS-evoked Ca^2+^ activation transiently contracted during days 1-4 but remained largely stable thereafter (Fig. 2E; Fig. S1Aii, Bii, and Cii). In contrast, visually evoked Ca^2+^ activation gradually expanded during the first week. Regarding temporal features, 25-Hz ICMS-induced activation duration initially shortened during the first few weeks, followed by a secondary increase during later stages (Fig. 2F; Fig. S1Aiii, Biii, Ciii), whereas 250-Hz ICMS-induced durations decreased monotonically. Visual stimulation, however, showed prolonged activation durations in chronic timepoints. Together, these trends suggest progressive tissue recovery and stabilization, where ICMS evokes spatially consistent yet temporally adaptive responses supporting reliable neuroprosthetic function.

Given the dynamic changes in Ca^2+^ activation over time, we next compared cross-condition relationships to determine whether the relative hierarchy among stimulation conditions remained stable chronically (Fig. S2). Across all timepoints, 25-Hz ICMS consistently evoked significantly stronger activation magnitudes than visual stimulation (post-hoc Tukey’s HSD test, *p* < 0.05, see Table S3 for detail), whereas visual stimuli typically produced the widest cortical spread except on day 0. Activation sizes for 25- and 250-Hz ICMS were comparable. Regarding response duration, although the rank order across stimulation conditions fluctuated, 25-Hz ICMS generally elicited longer-lasting activation than 250-Hz ICMS, except acutely on day 0. Overall, these results indicate that despite ongoing tissue remodeling, the relative spatial and temporal structure of ICMS-evoked Ca^2+^ activation remains stable, underscoring chronic functional biocompatibility and predictable network engagement over time.

### 3.2. Ca^2+^ depression induced by visual stimulation became smaller and shorter after chronic implantation

Next, we explored how the stimulation-evoked Ca^2+^ depression progressed over days following probe insertion (Fig. 3). Compared to Ca^2+^ activation, Ca^2+^ depression in this example mouse showed less consistent changes between day 0 and day 21 (Fig. 3A-C). Ca^2+^ depression induced by 25-Hz ICMS at day 21 was stronger and more spatially extensive than on day 0 (increase in blueish region in Fig. 3Aiv compared with ii). In contrast, Ca^2+^ depression induced by 250-Hz ICMS and visual stimulation at day 21 became weaker and/or spatially smaller than day 0.

Chronological analysis revealed that depression magnitudes remained relatively stable over time (Fig. 3D), although Ca^2+^ depression elicited by 250-Hz ICMS and visual stimulation tended toward reduction in chronic days (Fig. S3Ai, Bi, and Ci). Similarly, the spatial extent of Ca^2+^ depression progressively decreased over days, particularly under 250-Hz ICMS and visual stimulation (Fig. 3E; Fig. S3Aii, Bii, and Cii). Duration of ICMS-induced Ca^2+^ depression was largely stable, whereas visual stimulation induced depression became progressively shorter over chronic timepoints (Fig. 3F; Fig. S3Aiii, Biii, Ciii). These findings suggest that low-frequency ICMS preserves stable, functionally substantial depression, whereas high-frequency ICMS and visual stimulation showed partial declines in both magnitude and spatial extent, reflecting ongoing tissue adaptation and highlighting chronic device-tissue interactions relevant to long-term biocompatibility.

We next asked whether the relative hierarchy of Ca^2+^ depression across stimulation conditions remained stable over days (Fig. S4). Depression magnitudes induced by ICMS, especially 25 Hz, were consistently larger than those induced by visual stimulation. Depression distances remained comparable across conditions. Similarly, 25-Hz ICMS produced longer-lasting depression at chronic stages. Together, these results indicate that low-frequency ICMS reliably drives robust and sustained Ca^2+^ depression over chronic implantation, supporting predictable network modulation and translational relevance for neuroprosthetic applications.

### 3.3. Chronic recovery and fluctuation of Ca^+2^ activation and silencing in cells and neuropils

Given the progressive changes in widefield Ca^2+^ activation and depression, we next examined localized cellular responses near the stimulation site using chronic two-photon imaging (Fig. 4). Cellular-scale Ca^2+^ imaging was performed with two-photon microscopy. We successfully recorded chronic cellular activity within the same area around the implanted probe, which exhibited somatic and neuropil activation (Fig. 4A) and depression (Fig. S5).

Activation magnitudes of somatic Ca^2+^ induced by ICMS declined over days but recovered on specific chronic days (Fig. 4B; Fig. S6Ai and ii). Somatic activation magnitude induced by visual stimulation also fluctuated over days, with an overall slight increase (Fig. 4B; Fig. S6Aiii). The activation duration in the 25-Hz ICMS condition also showed non-monotonic changes over days (Fig. 4C; Fig. S6Bi), decreasing in the first several weeks, and then increasing during the chronic period. In contrast, activation duration under 250-Hz ICMS monotonically decreased over days (Fig. 4C; Fig. S6Bii), approaching the stimulation train duration. Visually-evoked somatic activation was consistently shorter than the stimulation duration but gradually lengthened over chronic days with fluctuations (Fig. 4C; Fig. S6Biii). These results suggest that visual stimulation produces transient responses independent of the implantation period, whereas ICMS, particularly at 25 Hz, can override transient neural dynamics, generating prolonged activation in specific neurons, a critical consideration for safe and effective neuroprosthetic parameter selection.

Although somatic depression magnitude was relatively stable compared to the somatic activation magnitude, there were still statistically significant fluctuations over days (LME model, *p* < 0.05; post-hoc Welch’s t-test with Holm-Bonferroni correction, *p* < 0.05; see Table S2 for detail), and the temporal dynamics differed across stimulation conditions (Fig. 4D, Fig. S6C). Depressions induced by 25-Hz ICMS and visual stimulation were relatively stable during the first 2-3 weeks, showed a transient increase, and then returned to or fell below initial magnitudes. In contrast, depression induced by 250-Hz ICMS monotonically increased over days. Somatic depression duration also varied significantly (Fig. 4E, LME model, *p* < 0.05, post-hoc Welch’s t-test with Holm-Bonferroni correction, *p* < 0.05, see Table S2 for detail; Fig. S6D, one-way ANOVA, *p* < 0.05, post-hoc Tukey’s HSD, *p* < 0.05, see Table S3 for detail). Apart from the longer duration induced by 25-Hz ICMS at day 0, depression generally lengthened between days 7 and 42. These patterns suggest that probe insertion transiently disrupts cellular depression mechanisms, which subsequently recover and reorganize, reflecting progressive tissue adaptation and establishment of functional biocompatibility.

Activation magnitude of excitatory Ca^2+^ in local neuropil induced by 25-Hz ICMS decreased several days after implantation, but recovered to day-0 levels (Fig. 4F, Fig. S6Ei). Also, the activation magnitude in neuropil induced by 250-Hz ICMS exhibited a similar trend (Fig. 4F; Fig. S6Eii). Together with somatic activation results, especially with 25-Hz ICMS, these observations suggest that implantation transiently desensitizes surrounding neural tissue to ICMS. Meanwhile, neuropil activation evoked by visual stimulation remained relatively constant across chronic days (Fig. 4F; Fig. S6Eiii). The activation duration in neuropil shortened initially under ICMS conditions, particularly at 25 Hz, and then lengthened over the chronic period (Fig. 4G; Fig. S6Fi and ii). Visual-stimulation-induced neuropil activation tended to lengthen in the first few days to approach stimulation duration (Fig. 4G; Fig. S6Fiii). Notably, several clusters under 25-Hz ICMS showed extremely prolonged durations exceeding 60 s, possibly due to some neurons or mice exhibiting extended activation (see 3.6). Neuropil depression magnitudes and durations fluctuated without significant trends (Fig. 4H-I; Fig. S6G-H), differing from somatic depression and suggesting distinct mechanisms govern somatic versus neuropil inhibitory dynamics.

Overall, these two-photon results indicate that ICMS-induced activation and depression undergo chronic modulation, reflecting the interplay between tissue adaptation, device-tissue interactions, and functional biocompatibility. The observed temporal and spatial recovery patterns are critical for optimizing long-term stimulation parameters and predicting neuroprosthetic performance.

### 3.4. Chronic hemodynamic OEF response gradually increased after probe insertion

Having observed progressive changes in neural activity at mesoscopic and cellular scales, we next investigated whether hemodynamic responses, measured by OEF, adapt over chronic timescales (Fig. 5). Evoked OEF reduction, reflecting relative oxygen supply increase, by 25-Hz ICMS spread through the cortex and persisted after stimulation offset on day 0, and was further enhanced by day 21 (Fig. 5A). Similarly, OEF reductions were enhanced on day 21 compared to day 0 in both 250-Hz ICMS and visual stimulation conditions (Fig. 5BC).

OEF magnitude progressively decreased over days (Fig. 5D; Fig. S7Ai, Bi and Ci), reflecting increased local metabolic supply likely associated with tissue recovery and adaptation after probe insertion. The spatial extent of OEF responses initially decreased during the first few days (Fig. E; Fig. S7Aii, Bii, and Cii), consistent with transient vascular disruption after implantation, and subsequently increased over a wider cortical area, likely reflecting reformation of vessel patterns similar to post-injury cortical recovery^86^. Response duration shortened during the initial weeks and then lengthened with the 25-Hz ICMS condition (Fig. 5F; Fig. S7Aiii, Biii, and Ciii), paralleling trends in mesoscopic-scale neural activation.

Across stimulation conditions, 25-Hz ICMS induced greater OEF reductions, particularly at later chronic stages (Fig. S8A), whereas visual stimulation produced the largest spatial extent (Fig. S8B), likely reflecting the full-field nature of visual input. ICMS responses also displayed longer durations than visual stimulation (Fig. S8C), exceeding the 10-s stimulation period, suggesting enhanced metabolic demand under artificial stimulation that may diverge from natural sensory processing.

These results indicate that ICMS chronically recruits oxygen supply in a frequency-dependent manner, revealing functional consequences of electrode-tissue interactions and highlighting considerations for safe and effective neuroprosthetic design. The gradual recovery and enhancement of hemodynamic responses support the functional biocompatibility of chronic ICMS devices, as local metabolism adapts to maintain network function over time.

### 3.5. Chronic probe insertion injury leads to spontaneous silencing

Beyond stimulation-induced silencing (Fig. 3), we quantified how probe insertion itself modulates spontaneous neural and hemodynamic activity (Fig. 6). Coefficient of variation (CV; standard deviation normalized to its mean, σ/μ) was used as a metric of tissue excitability, where larger values indicate greater fluctuations in signals, reflecting more neural and hemodynamic activity. During pre-stimulation periods, CV was slightly reduced around the probe at early time points (day 0), while medial and lateral cortical regions near craniotomy edges exhibited lower CV, likely due to collateral tissue damage from surgical manipulation (Fig. 6A). By day 14, global CV increased, particularly for spontaneous neural activity (Fig. 6B, top), indicating gradual recovery and network adaptation.

To quantify localized silencing, we computed SI_350_ normalized to CV at 300 to 400 µm from the probe (SI_350_, (CV-CV_350_)/CV_350_; Fig. 6B, bottom). Early post-implantation SI_350_ values near the probe were significantly below zero for both neural and hemodynamic activity (one-sample t-test with Bonferroni correction, *p* < 0.05, see Table S2 for details), indicating enhanced local silencing proximal to the electrode (Fig. 6B, bottom). Similar trends were observed during visual stimulation (Fig. 6C, D). These findings indicate that probe insertion transiently suppresses local spontaneous activity, with effects persisting for several weeks. The dynamic recovery of activity over chronic time points reflects functional biocompatibility, as local networks regain excitability while adapting to the implanted device. Understanding these localized silencing patterns is critical for optimizing long-term ICMS performance and safety in translational neuroprosthetic applications.

### 3.6. Chronic 25-Hz ICMS preferentially induces epileptiform neural activity

Mesoscopic- and microscopic-scale recordings revealed that ICMS occasionally elicited extremely strong and widespread neural activation, consistent with putative epileptiform events. Epileptiform neural activity reflects synchronized, pathological firing of a subset of neurons, which is distinct from cortical spreading depolarizations, a slow, propagating wave of near-complete depolarization affecting all nearby neurons and glia, followed by a transient activity depression. Such events represent a critical safety consideration for chronic neuroprosthetic applications, as excessive or prolonged activation can compromise functional biocompatibility and long-term device performance. An example epileptiform event exhibited large-magnitude activation propagating across the entire visual cortex and persisting >10 s after stimulation offset (Fig. 7A, Mov. S1), in contrast to non-epileptiform activity in the same animal (Mov. S2). These events occurred almost exclusively during 25-Hz ICMS (Fig. 7B), indicating a frequency-dependent susceptibility.

**Figure 7.**
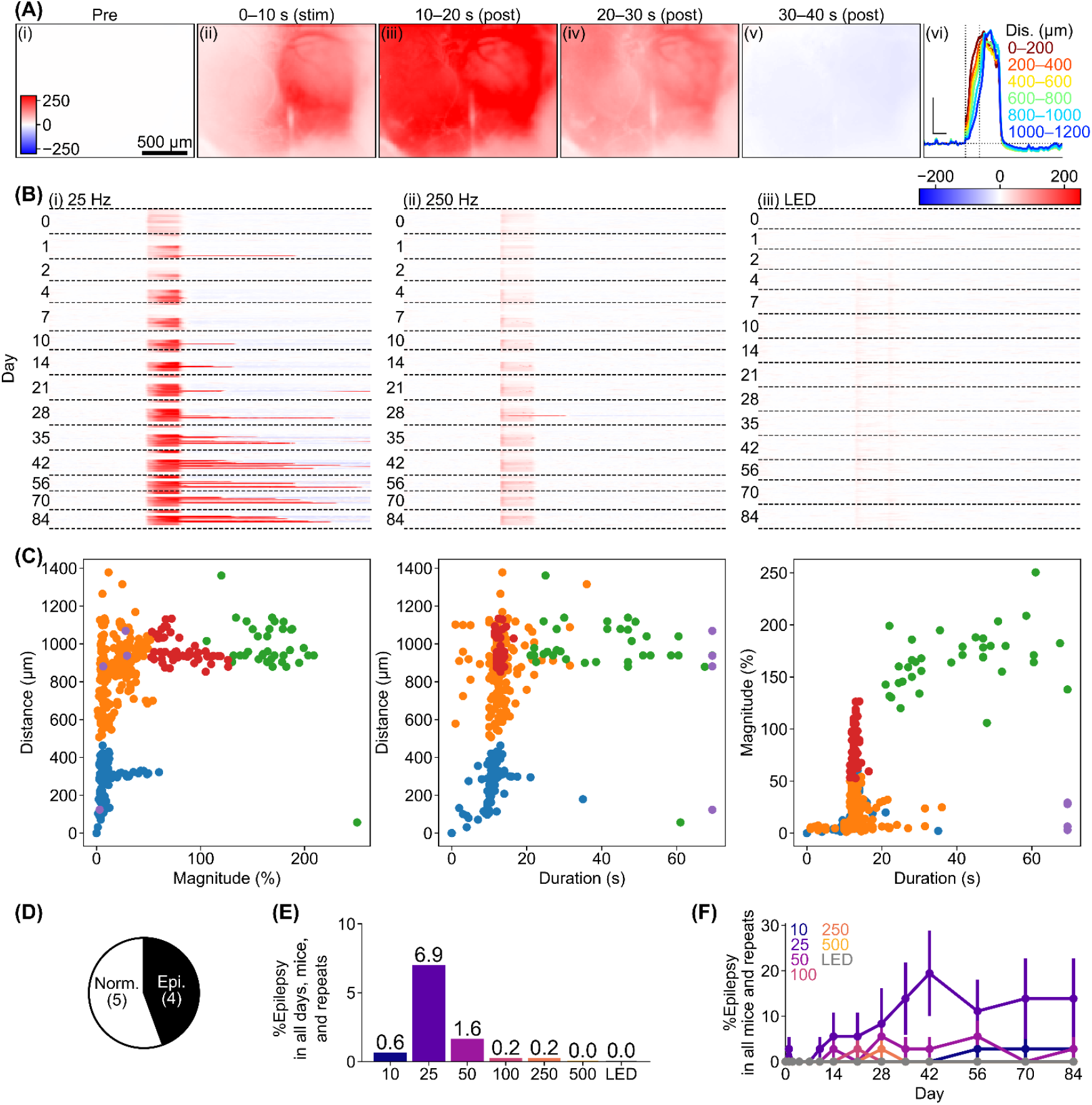
Chronic emergence of epileptiform Ca^2+^ activity is preferentially induced by 25-Hz ICMS. (**A**) Example epileptiform Ca^2+^ activity induced by 25-Hz ICMS, showing extremely strong activation spreading across the entire visual cortex and persisting >10 s after stimulation offset (i-v, maps; vi, time courses at each distance). (**B**) Overview of Ca^2+^ activity induced by 25-Hz ICMS (i), 250-Hz ICMS (ii), and visual stimulation (iii). Each line represents one mouse. (**C**) Distributions of magnitude, duration, and spatial spread of Ca^2+^ activation. Colors indicate clusters identified using agglomerative clustering (see Methods). Epileptiform activity was defined from the green cluster with magnitude >100%, duration >20 s, and distance >800 µm. (**D**) Number of mice exhibiting epileptiform activity at least once across all days, conditions, and repeats (Epi) versus not (Norm). (**E**) Probability of epileptiform activity across stimulation conditions for all days, mice, and repeats. (**F**) Probability of epileptiform activity over days for all ICMS and visual stimulation conditions. Mean ± SEM across animals.

Agglomerative clustering of activation magnitude, duration, and spatial spread revealed a distinct cluster representing epileptiform activity (magnitude >100%, duration >20 s, distance >800 µm; Fig. 7C, green; Fig. S9). Using these criteria, 4 out of 9 mice exhibited epileptiform events at least once across repeats, stimulation conditions, and post-implantation days (Fig. 7D). The 25-Hz ICMS condition strongly entrained epileptiform activity compared with all other tested stimulation parameters (Fig. 7E; Fig. S10). The occurrence rate of epileptiform events increased gradually over chronic implantation days, peaking around day 42 (∼15% of trials), and then plateaued (Fig. 7F; Fig. S11). Event magnitude remained largely stable over time (Fig. S12), indicating persistent network susceptibility despite tissue adaptation, highlighting the importance of chronic biocompatibility assessment. Lower (10-Hz) and higher (50-250 Hz) ICMS frequencies occasionally elicited epileptiform activity in mid-to late-chronic periods but at lower prevalence. These results indicate that 25-Hz ICMS presents a heightened risk for inducing epileptiform neural activity in chronically implanted cortex, which must be carefully considered when designing safe, long-term neuroprosthetic stimulation protocols.

### 3.7. Neuro-vascular coupling progressively recovers after probe insertion

Given that both the neural and hemodynamic signals exhibited characteristic recovery processes, we next assessed the functional coupling between these two signals (Fig. 8; Fig. S13 for calculation, see 2.4.6 for details). By calculating the deconvolution of time courses of neural and hemodynamic signals (Fig. 8A), we obtained hemodynamic response function (HRF; Fig. 8B) that indicates the dynamic scaling factor converting neural activity into vascular responses, providing a functional measure of interface biocompatibility.

**Figure 8.**
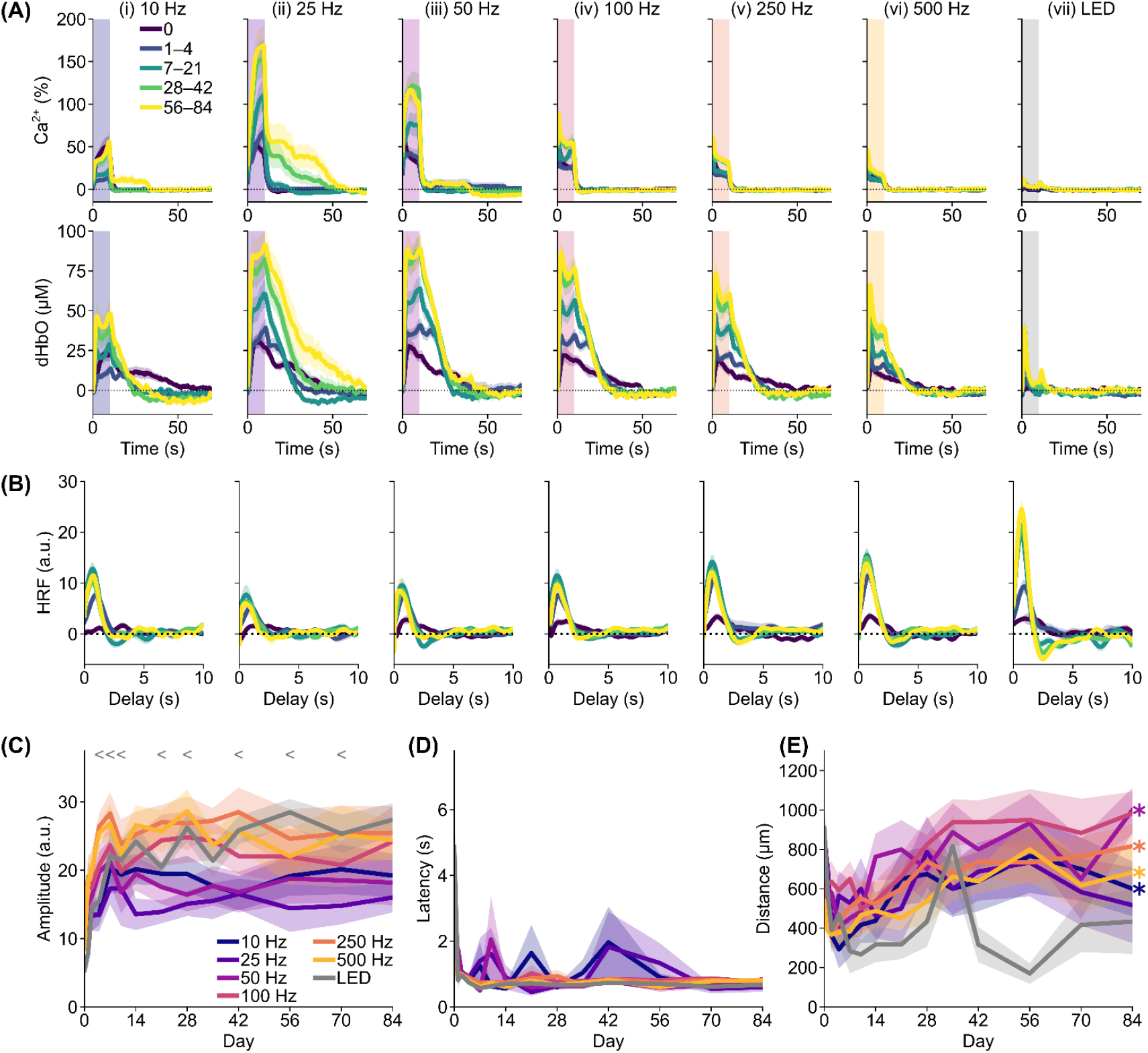
Neurovascular coupling progressively recovers after probe insertion. (**A**) Ca^2+^ and OEF time courses around the electrodes (< 100 μm, Mean ± SEM across mice). (**B**) Corresponding hemodynamic response function (HRF) profiles. (**C-E**) Comparisons of HRF peak amplitude (C), peak latency (D), and peak distance (E) across days after probe insertion. Mean ± SEM across mice; ^*^ next to line plot indicates significant effects of day (LME model), “<“ and “>“ indicate significant differences compared with days 0 and 84, respectively (LME model followed by paired t-test with Holm-Bonferroni correction).

On day 0, ICMS-induced HRFs were generally small (Fig. 8Bi), indicating weak coupling between neural activity and vascular response immediately after probe insertion. By days 7-21, HRF amplitudes increased across all ICMS frequencies, corresponding to greater vascular responsiveness for a given neural signal. Peak latencies shortened over this period, demonstrating faster and stronger neurovascular coupling with chronic adaptation. Visual stimulation elicited smaller neural signals but relatively larger hemodynamic responses, producing higher HRFs, consistent with intact subcortical visual pathways maintaining stable coupling. A post-peak trough was evident under visual stimulation, possibly reflecting balanced excitatory and inhibitory network entrainment, whereas this feature was less apparent for ICMS, indicating distinct or less coordinated recruitment of inhibitory circuits. We also obtained qualitatively similar results for HRF reconstructed from Ca^2+^ and OEF (Fig. S14).

Spatial analysis revealed HRF amplitude was reduced near the probe, particularly under chronic ICMS conditions (Fig. S15), suggesting persistent local perturbation and incomplete tissue recovery (Fig. 6). Peak HRF locations progressively shifted away from the probe, indicating vascular network reorganization and distal functional compensation during chronic recovery. In contrast, visual stimulation showed minimal local weakening, highlighting preservation of natural neurovascular signaling in uninjured pathways.

Statistical quantification confirmed that HRF amplitudes at day 0 were significantly lower than those at later time points (Fig. 8C, Fig. S16A, one-way ANOVA followed by post-hoc Tukey’s HSD test, *p* < 0.05, see Table S3 for details), while peak latencies decreased progressively (Fig. 8D, Fig. S16B). Under visual stimulation, HRF latency distributions narrowed over chronic periods, producing higher average peak values, indicating stabilization of neurovascular responses. Most prominent changes occurred within the first few days to weeks, defining a critical window for early tissue recovery and biocompatibility stabilization.

Peak HRF distances initially decreased or remained stable until days 1-21, then expanded chronically for both ICMS and visual stimulation conditions (Fig. 8E, Fig. S16C). This chronic enlargement of responsive regions reflects distal vascular remodeling, supporting sustained neurovascular coupling and overall interface performance. Together, these results demonstrate that probe insertion acutely disrupts neurovascular function, but chronic recovery leads to progressive restoration of coupling, spatial reorganization, and stabilization of vascular responsiveness (Fig. 5D-F; Fig. S7Aii, Bii, and Cii), providing a framework to optimize safe, long-term ICMS parameters for translational neuroprosthetic applications.

## 4.0 Discussion

This study longitudinally characterized mesoscale and cellular network responses to ICMS and visual stimulation in mouse visual cortex for 12 weeks following microelectrode implantation. Results showed frequency- and time-dependent remodeling of excitability, hemodynamics, and neurovascular coupling, with progressive changes in activation magnitude, spatial spread, and temporal dynamics. Among tested conditions, 25-Hz ICMS consistently posed the highest risk for late-onset, long-duration epileptiform activity, underscoring a key safety consideration for chronic neuroprosthetic use.

### 4.1 Acute neural activity suppression after intracortical probe insertion

In this study, we evaluated the chronic alteration of cortical Ca^2+^ activity in awake mice induced by two principal frequencies of ICMS (Figs. 2-4) and visual stimulation (Fig. 6C-D). Some properties gradually changed over days in the chronic period (Fig. 2), whereas other properties, such as somatic activation magnitudes under two-photon microscopy (Fig. 4), exhibited marked shifts between day 0 and days 1-4. This acute alteration in neural activity (Fig. 6) likely reflects both mechanical injury from probe insertion and residual effects of anesthesia.

Mechanical injury from probe insertion triggers a cascade of acute tissue responses beginning with direct physical damage such as neuronal strain and membrane rupture, which compromise cellular integrity and excitability^31,52^. Concurrently, blood vessel rupture occurs^50,87^, causing localized hemorrhage and disruption of the blood-brain barrier, which exacerbating inflammatory signaling^24,65^. Vascular damage also induces pericyte constriction^38^, further compromising local blood flow and oxygen delivery and contributing to acute ischemia^1^. The injury activates resident microglia^64^, which adopt a pro-inflammatory phenotype, and induces astrocyte hypertrophy^54^, both hallmarks of reactive gliosis that modulate the local microenvironment. Oligodendrocyte precursor cells (OPCs) respond by migrating^88^, while demyelination occurs in nearby axons^51^, impairing conduction and network function^26^. These combined effects attract peripheral immune cells, notably neutrophils and monocytes^1,58^, which infiltrate the injury site and produce reactive oxygen species (ROS)^89^. Elevated ROS

levels exacerbate oxidative stress, damaging neuronal membranes, proteins, and DNA, and further impairing metabolic function. These combined effects of vascular compromise reducing metabolic supply^1,50^, inflammation increasing metabolic consumption^73^, and oxidative stress^33^ create an ischemic microenvironment that impairs neural activity (including inhibitory neurons)^26,52,90,91^, network stability^26^, and silencing sensory-evoked local neural activity^1,45^ (Fig. 6C and D). These injury-induced processes collectively contribute to the observed decrease in Ca^2+^ activity by disrupting neuronal integrity, synaptic function, and the neurovascular support critical for cortical excitability.

Day 0 responses likely reflect the combined effects of acute insertion injury and residual anesthesia. Because ICMS without probe insertion is not feasible, the influence of anesthesia warrants particular consideration. The ketamine-xylazine mixture used before surgery, while standard for rodents, induces lasting depression of neural activity and cerebrovascular tone beyond its sedative window^92^. Although imaging began ∼2 h after induction, residual hypothermia, hypoxia, and suppressed synaptic activity may have persisted. The gradual recovery of ICMS-evoked responses from day 0 to days 1-4 likely reflects the combined resolution of anesthetic effects and early tissue stabilization^93,94^. Future studies employing sham operations and alternative anesthetic regimens will be essential to disentangle anesthesia-related suppression from insertion-induced perturbations.

### 4.2 Chronic increased activation magnitude correlates with epileptiform activity across frequencies, with heightened risk at 25 Hz

We also observed epileptiform cortical Ca^2+^ activity in a subset of mice, particularly with 25-Hz ICMS during late chronic days (Fig. 7). Previous studies have linked epilepsy to traumatic brain injury (TBI)^95^. Because this study used a single genetic strain (Thy1-GCaMP6s), genetic factors are unlikely to explain the selective occurrence of epileptiform activity. This activity, along with increased mesoscale activation magnitude, may relate to a decreased inhibitory neuron activity^26^, while excitatory neurons remained stable over chronic days post-implantation. These stimulation-evoked after-discharges reflect persistent network activity following stimulus offset rather than cortical spreading depolarization^1,52^ (characterized by a slow propagating wave depolarization of every neuron), and they appear on a timescale and spatial profile consistent with stimulation-entrained hyperexcitability (Fig. 7).

Clinically, late post-traumatic seizures, occurring more than one week after injury, are associated with intracranial hematoma, depressed skull fracture, and dural penetration^96^, conditions analogous to our surgical procedures. Following TBI timelines^97,98^, inflammation and gliosis around the implant develop over days to weeks^89,99,100^, processes believed to initiate epileptogenesis^95^. Anti-inflammatory treatment (e.g.dexamethasone^63^) may mitigate epileptiform risk. However, it is also important to consider that corticosteroids increase mortality in TBI by suppressing the immune system, raising infection risk, inducing hyperglycemia, impairing tissue repair, and exacerbating complications^101^.

For neuroprosthetic applications, 25-Hz stimulation effectively drives neural activity but may require careful adjustment of parameters such as amplitude, pulse duration, and number of simultaneously stimulated channels to reduce seizure risk. Other studies have reported that after-discharges following cortical stimulation can persist independently, often increasing seizure risk, with focal aware seizures observed despite safety measures^102^. This study further showed that not all electrodes equally contribute to seizure-like activity. Similarly, our results showed that only a subset of animals and stimulation electrodes exhibiting these events (Fig. 7D-F), indicating electrode- and state-dependent susceptibility. Continuous EEG monitoring may facilitate early detection and timely termination of stimulation. While antiseizure medications like phenytoin may reduce ICMS-triggered epileptiform activity, efficacy is limited to early seizures. Future studies with larger populations could identify non-genetic factors distinguishing epileptiform from non-epileptiform responses, including potential alterations in inhibitory neuron populations and function, thereby informing safer ICMS strategies. Closed-loop neural engineering approaches combining ICMS with electrophysiological recording may optimize 25-Hz stimulation by aborting or adjusting stimulation upon detecting epileptiform activity^103^. Additionally, addressing biomaterials challenges, such as minimizing tissue disruption and chronic inflammatory responses around implanted electrodes, remains critical to ensure long-term functional stability and biocompatibility.

### 4.3 Chronic probe injury yields partial recovery but progressively disrupts local neurovascular coupling and neuronal activity despite global improvement

Over weeks following implantation, both neural and hemodynamic measures partially recover (Figs. 2, 5, 6). At the mesoscale, Ca^2+^ activation magnitudes increased with days post-insertion (Fig. 2), with 25-Hz ICMS producing progressively larger Ca^2+^ responses and visually evoked Ca^2+^ responses strengthening over time. At the cellular scale (Fig. 4), two-photon imaging showed recovery of activation silencing in soma and neuropil, and population power increased toward patterns consistent with awake GCaMP activity (Fig. 6), reflecting recovery of baseline excitability within the first 28 days. Hemodynamically, OEF magnitudes progressively decreased from baseline (Fig. 5), consistent with a relative increase in oxygen supply accompaning tissue recovery.

The HRF became larger in amplitude and faster in latency across days independent of stimulation condition (Figs. 8A, C, D), suggesting either improved spontaneous neurovascular coupling or relatively greater restoration of baseline hemodynamic function versus neural activity. Because HRF improvements were near-independent of stimulation parameters, frequency-specific hyperexcitability and epileptiform events are unlikely to reflect a global mismatch between neural demand and vascular supply. Spontaneous-activity metrics nuance this interpretation. Coefficient of variation for neural signals increased from day 0 to day 14, indicating recovery from acute silencing rather than a paradoxical decline in neural activity. Therefore, HRF enhancement cannot be attributed solely to disproportionately improved baseline hemodynamics. Nonetheless, HRF amplitude was consistently attenuated in the immediate vicinity of the probe (Fig. S15), revealing a focal and persistent deficit in neurovascular coupling^104^.

Multiple mechanistic pathways sustain this local deficit as discussed in Section 4.1 or relevant reviews^1,24,65,105^, including the remodeling of microcirculation and capillary architecture near the implant^1,66^, perfusion capacity (Figs. 5 and 6), and impairing clearance of metabolic waste^56,57^. Implant-induced damage to myelin and oligodendrocytes^27,28,51,74,91,106,107^ compromises axonal conduction and increases the metabolic cost of action potential propagation. At the same time, insufficient remyelination leaves axons vulnerable to further degeneration. In parallel, chronic dysregulation of the autophagy-lysosomal pathway^56^ impairs degradation of protein aggregates and damaged organelles^57^, exacerbating oxidative stress and inflammatory injury. Collectively, these vascular, myelin, and clearance deficits likely maintain the ischemic and neurotoxic microenvironment near the probe, underlying the persistent local neurovascular uncoupling (Fig. 8) and reduced neuronal calcium activity (Fig. 6), despite evidence of global recovery in hemodynamic and neural responses.

Despite focal deficits, HRF peak latencies in chronic imaging approximate values previously reported in awake mice^75^, indicating substantial recovery of neurovascular timing at larger scales. Notably, Fig. 6 shows that neuronal calcium activity recovers in parallel with OEF recovery, suggesting that improved oxygen delivery supports restoration of neuronal function. However, persistent local impairments, such as attenuated HRF amplitudes and sustained local silencing indicate incomplete biocompatibility at the electrode-tissue interface. These focal impairments arise from mechanical injury^31,51-53,108-110^, disrupted microcirculation^1,38,40,50,65^, chronic inflammation^29,54,59,61,69,111-113^, and accumulation of metabolic waste^56,57^, which collectively create a local neurotoxic environment and contribute to long-term interface functional degradation. Addressing these challenges requires next-generation biomaterials designed to minimize tissue disruption, preserve vascular integrity, and mitigate ischemia and neurotoxicity^105^. Optimized surface chemistries and mechanically compliant coatings can reduce inflammation, promote glial and vascular health, and facilitate metabolic support, thereby sustaining neurovascular coupling and neuronal activity over chronic implantation periods^62,114-116^. Such material innovations are critical for improving long-term ICMS performance, stability, and overall functional biocompatibility^24^.

### 4.4 Limitation and Future Directions

This study establishes chronic, frequency-dependent remodeling of cortical excitability and perfusion after microelectrode implantation, but constraints limit mechanistic inference and generalizability. First, the imaging platform reports primarily excitatory-cell calcium (Thy1-GCaMP6s), which does not resolve inhibitory subclasses^35,37,117,118^, astrocytes^54^, oligodendrocytes^51^, OPCs^88^, microglia^61-64,66,115,119,120^, or pericytes^38^ that critically shape biocompatibility of ICMS electrodes over time. Second, electrode insertion produces focal vascular rupture, protein deposition, and blood-derived inflammatory signals that progress with pericyte-driven vascular remodeling over weeks. These vascular and blood-brain barrier effects can directly alter OEF, neurovascular coupling^121,122^, and local excitability^50,87,109,111,123^, thereby influencing the functional biocompatibility of ICMS (the ability to sustain effective network activation with acceptable tissue responses^25^). Third, calcium and widefield OEF are indirect proxies for spiking and metabolic flux, leaving the cellular drivers of epileptiform-like events, the role of local field dynamics, and the precise timing of excitation versus inhibition ambiguous^124^. While voltage indicators are available for *ex vivo* slice studies, their signal-to-noise ratio remains insufficient for chronic in vivo experiments with chronically implanted intracortical microelectrodes^125^. Finally, the etiology of 25-Hz linked, chronic epileptiform events is unresolved^126^. Frequency-dependent entrainment and state-dependent susceptibility are plausible^127,128^, but causal validation is required. Future work should integrate multi-cell-type imaging, high-resolution electrophysiology, and metabolic assays to quantify how foreign body responses alter long-term functional biocompatibility of stimulation interfaces. Despite limitations, these findings provide physiological constraints and safety-relevant biomarkers to guide the design of safer, longer-lasting cortical neuroprostheses.

## Conclusion

We characterized chronic changes in excitatory neuronal activity and local hemodynamics following implantation of four-channel microelectrode arrays in mouse V1, using widefield calcium imaging, two-photon somatic and neuropil imaging, and OEF mapping over 0-84 days. Probe insertion acutely caused focal suppression of spontaneous and evoked activity, highlighting the initial impact on tissue biocompatibility. Over ensuing weeks, ICMS-evoked Ca^2+^ responses increased in magnitude while spatial spread remained largely stable, with temporal dynamics exhibiting frequency dependence. Hemodynamic responses and neurovascular coupling progressively recovered, reflecting restoration of vascular function and tissue compatibility. Notably, chronic 25-Hz ICMS preferentially induced large, spreading, long-lasting epileptiform-like events, emphasizing the need to carefully tailor stimulation parameters. These results also underscore the critical role of biomaterials in mitigating local tissue injury, preserving vascular integrity, and sustaining neuronal and hemodynamic function, pointing to the need for next-generation electrode designs with improved surface chemistry, mechanical compliance, and biocompatibility. Taken together, our findings define physiological constraints for chronic cortical stimulation and provide mechanistic guidance for optimizing stimulation protocols and microelectrode design, advancing safe, durable, and translationally relevant neural interfaces.

## Acknowledgement

This work was supported by: NIH R01NS105691, NIH R01NS115707, NIH R03AG072218, NIH R01NS129632 and NSF CBET CAREER 1943906, Senior Vice-Chancellor’s Pitt Momentum Grant, PIT-NI.

