## SupplementalFigures for "Chronic alteration of Ca^2+^ and hemodynamic signals induced by intracortical microstimulation in the visual cortex of awake mice"

### Supplementary figures

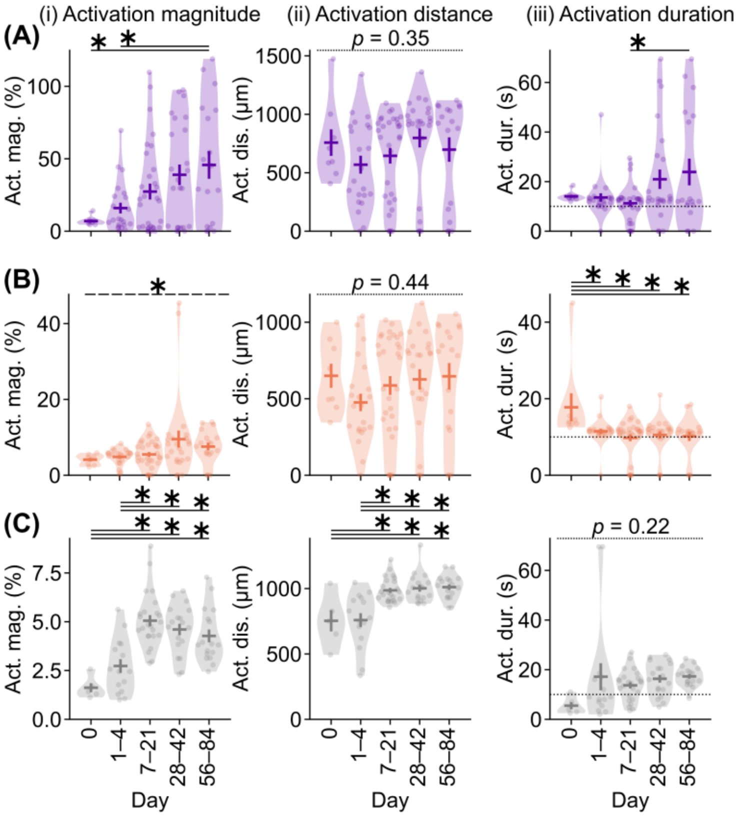


**Figure S1. Progressive recovery and stimulation-specific dynamics of cortical $Ca^{2+}$ responses following probe implantation.** Comparisons of $Ca^{2+}$ activation (i) magnitude, (ii) spatial spread, and (iii) duration under (A) 25-Hz ICMS, (B) 250-Hz ICMS, and (C) visual stimulation conditions across days after probe insertion. The horizontal dotted line in (Aiii), (Biii), and (Ciii) marks the 10-s stimulation period. Horizontal and vertical bars represent mean ± SEM across samples. * with solid lines denote $p < 0.05$ for post-hoc Tukey's HSD pairwise comparisons; * with dashed lines indicate $p < 0.05$ for one-way ANOVA without significant post-hoc differences; "*p*" with dotted lines indicates nonsignificant one-way ANOVA results.

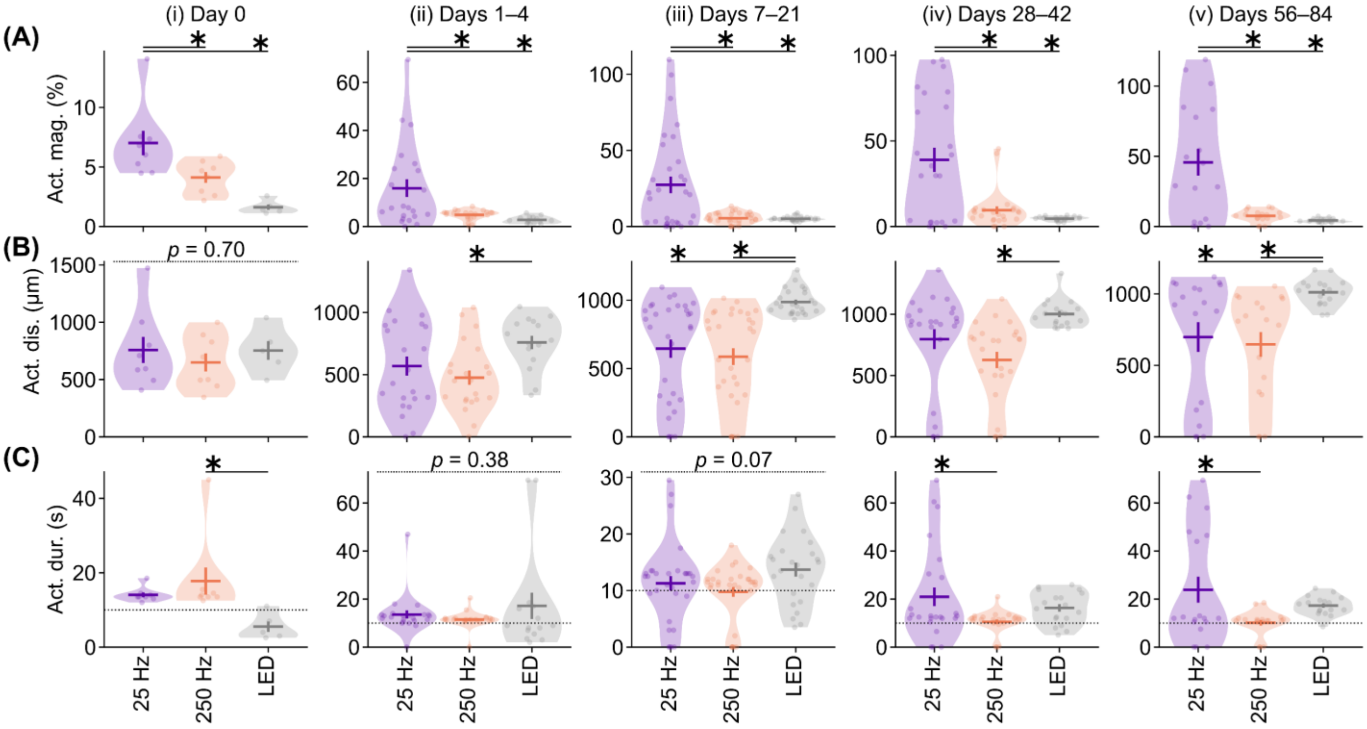


**Figure S2. Temporal evolution of ICMS-evoked cortical $Ca^{2+}$ dynamics across chronic implantation.** Comparisons of $Ca^{2+}$ activation (A) magnitude, (B) spatial spread, and (C) duration at (i) day 0, (ii) days 1–4, (iii) days 7–21, (iv) days 28–42, and (v) days 56–84 across stimulation conditions. The horizontal dotted line in (C) denotes the 10-s stimulation period. Horizontal and vertical bars represent mean ± SEM across samples. * with solid lines indicate $p < 0.05$ for post-hoc Tukey's HSD pairwise comparisons; "$p$" with dotted lines denotes nonsignificant one-way ANOVA results.

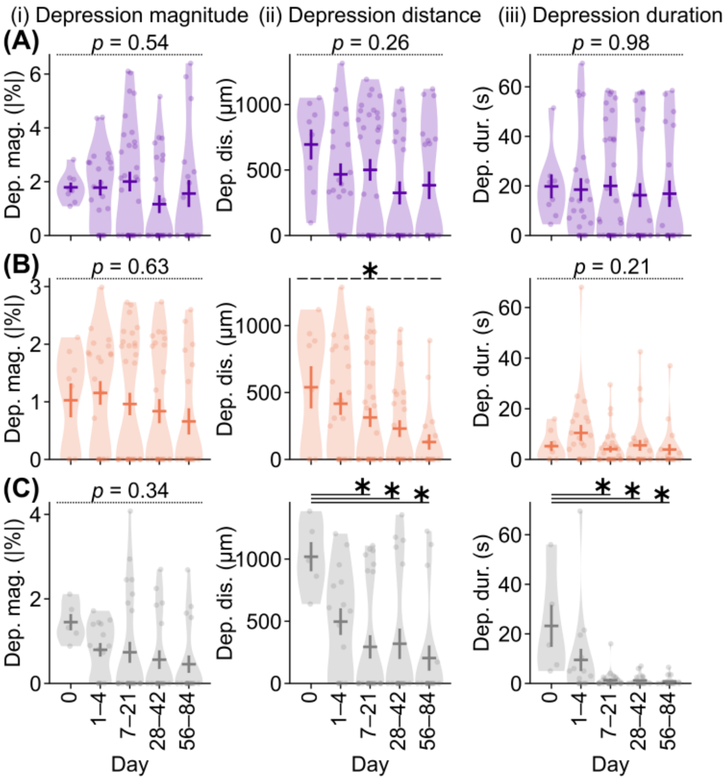


**Figure S3. Chronic suppression of cortical $Ca^{2+}$ activity reveals long-term inhibitory effects of ICMS and implantation.** Comparisons of $Ca^{2+}$ signal depression (i) magnitude, (ii) spatial extent, and (iii) duration under (A) 25-Hz ICMS, (B) 250-Hz ICMS, and (C) visual stimulation across days after probe insertion. Horizontal and vertical bars represent mean ± SEM across samples. * with solid lines indicate $p < 0.05$ for post-hoc Tukey's HSD pairwise comparisons; * with dashed lines indicate $p < 0.05$ for one-way ANOVA without post-hoc significant differences; "$p$" with dotted lines denotes nonsignificant one-way ANOVA results.

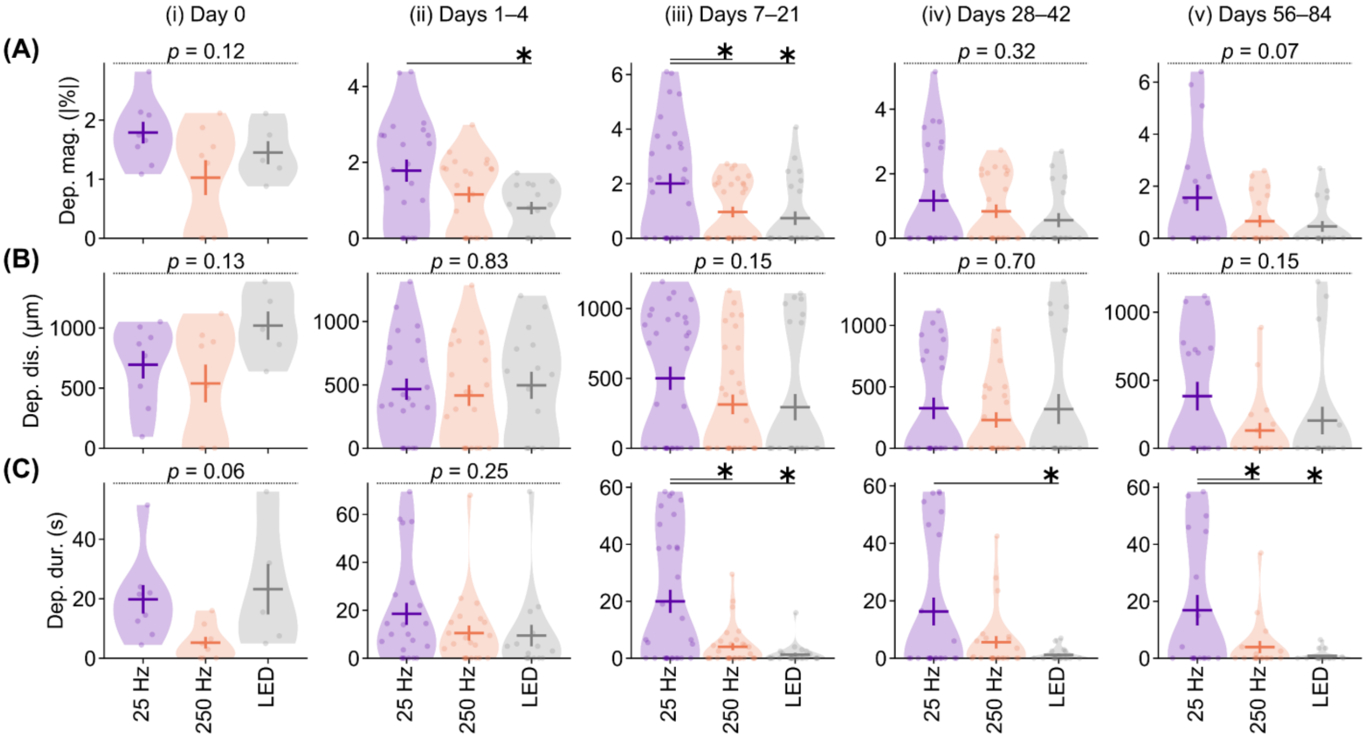


**Figure S4. Temporal evolution of cortical $Ca^{2+}$ suppression reveals partial recovery of excitatory signaling over chronic implantation.** Comparisons of $Ca^{2+}$ signal depression (A) magnitude, (B) spatial extent, and (C) duration at (i) day 0, (ii) days 1–4, (iii) days 7–21, (iv) days 28–42, and (v) days 56–84 across stimulation conditions. Horizontal and vertical bars represent mean ± SEM across samples. * with solid lines indicate $p < 0.05$ for post-hoc Tukey's HSD pairwise comparisons. "*p*" with dotted lines denotes nonsignificant one-way ANOVA results.

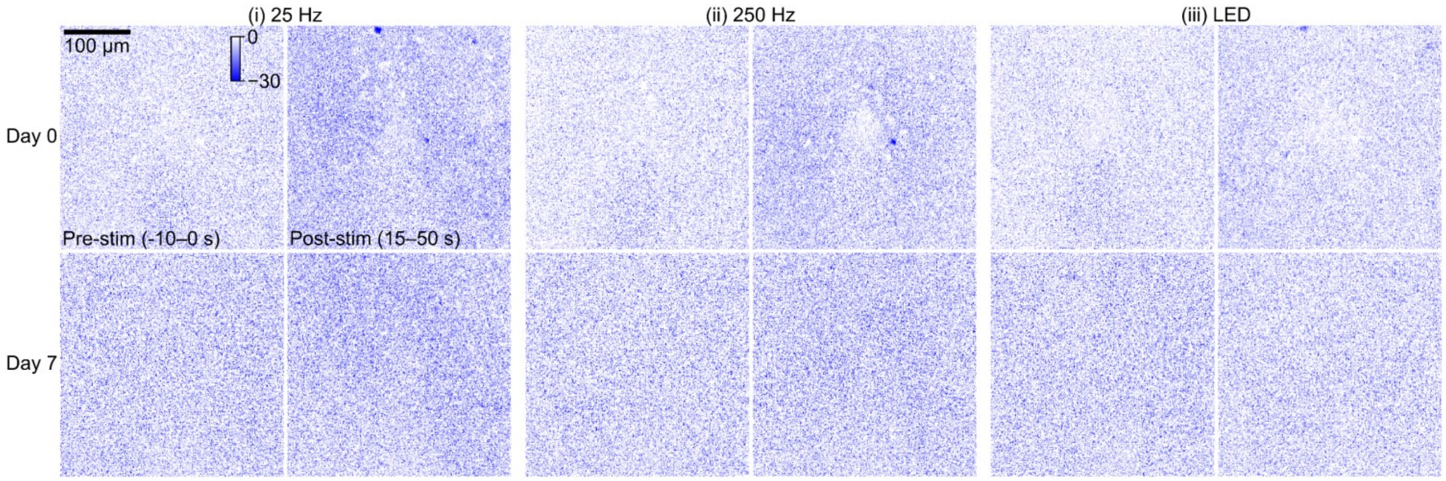


**Figure S5. Two-photon imaging reveals transient $Ca^{2+}$ suppression near implant that diminishes over the first week post-insertion.** Representative two-photon $Ca^{2+}$ images at day 0 (top) and day 7 (bottom) during (i) 25-Hz ICMS, (ii) 250-Hz ICMS, and (iii) visual stimulation. Blue indicates decreases in $Ca^{2+}$ signal relative to pre-stimulation baseline. Images are from the same mouse shown in Fig. 4A.

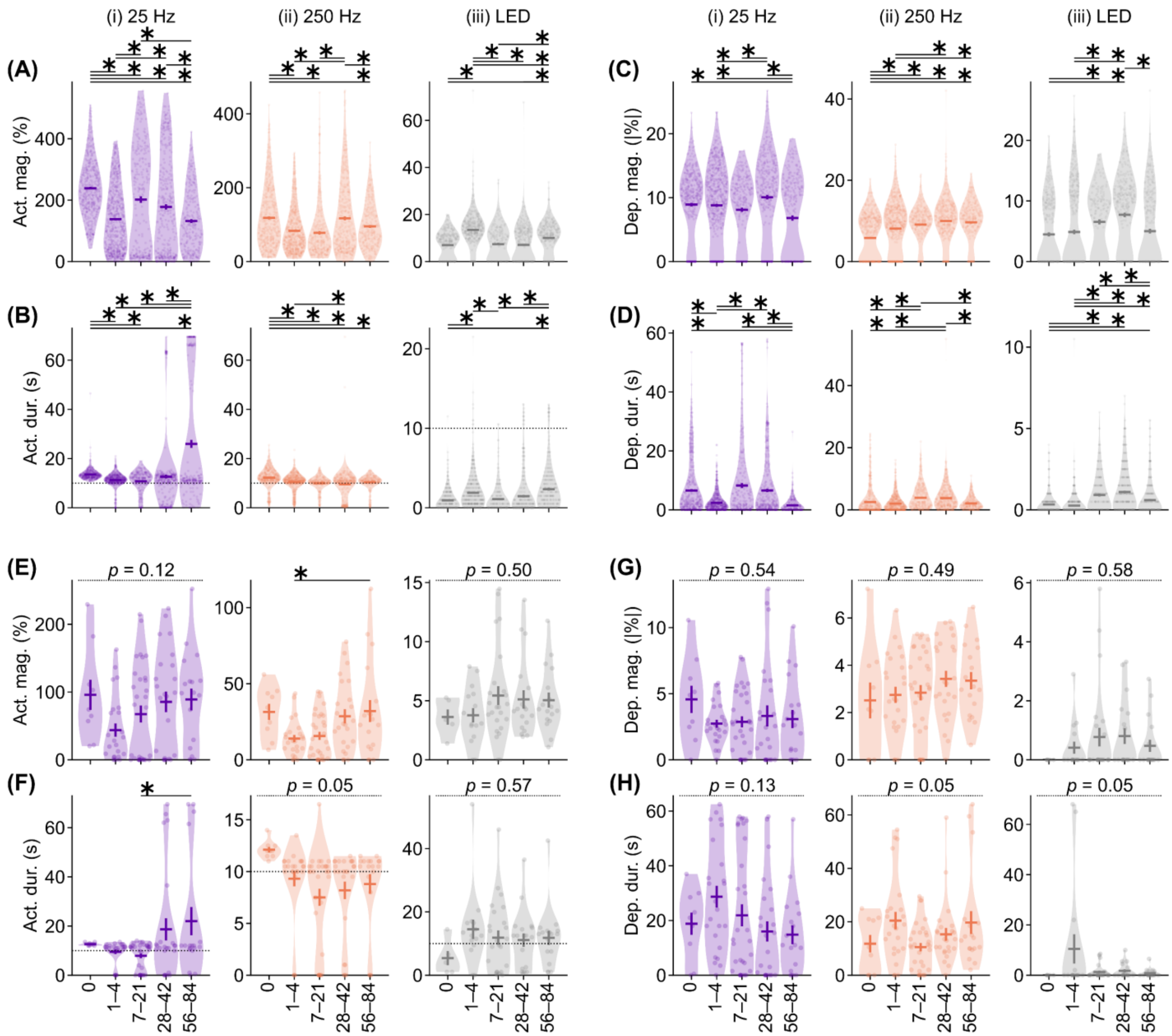


**Figure S6. Differential recovery of somatic and neuropil $Ca^{2+}$ responses reveals compartment-specific adaptation after chronic implantation.** (A–B) Somatic activation (A) magnitude and (B) duration under (i) 25-Hz ICMS, (ii) 250-Hz ICMS, and (iii) visual stimulation across days post-implantation. (C–D) Somatic depression (C) magnitude and (D) duration across days. (E–F) Neuropil activation (E) magnitude and (F) duration across days. (G–H) Neuropil depression (G) magnitude and (H) duration across days. Symbols represent individual neurons (A–D) or mice (E–H). Horizontal and vertical lines indicate mean ± SEM. Horizontal dotted lines in (B) and (F) mark stimulation duration (10 s). * with solid line denotes $p < 0.05$ for post-hoc Tukey's HSD pairwise comparisons. $p$ with dotted line indicates $p$-value for one-way ANOVA without significant difference.

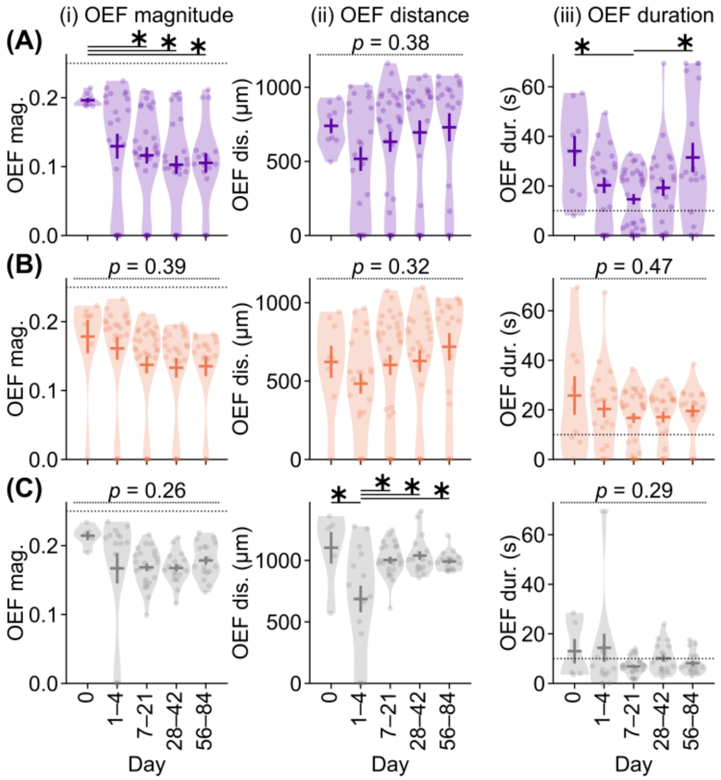


**Figure S7. Hemodynamic recovery dynamics show progressive normalization of oxygen extraction following probe insertion.** (A–C) Comparisons of OEF (i) magnitude, (ii) spatial extent (distance), and (iii) duration under (A) 25-Hz ICMS, (B) 250-Hz ICMS, and (C) visual stimulation across days post-implantation. Horizontal dotted lines in (Ai), (Bi), and (Ci) denote the pre-stimulation baseline OEF (0.25). Horizontal dotted lines in (Aiii), (Biii), and (Ciii) indicate stimulation duration (10 s). Horizontal and vertical lines represent mean ± SEM across samples. * with solid line indicates $p < 0.05$ for post-hoc Tukey's HSD pairwise comparisons. $p$ with dotted line indicates $p$-value for one-way ANOVA without significant difference.

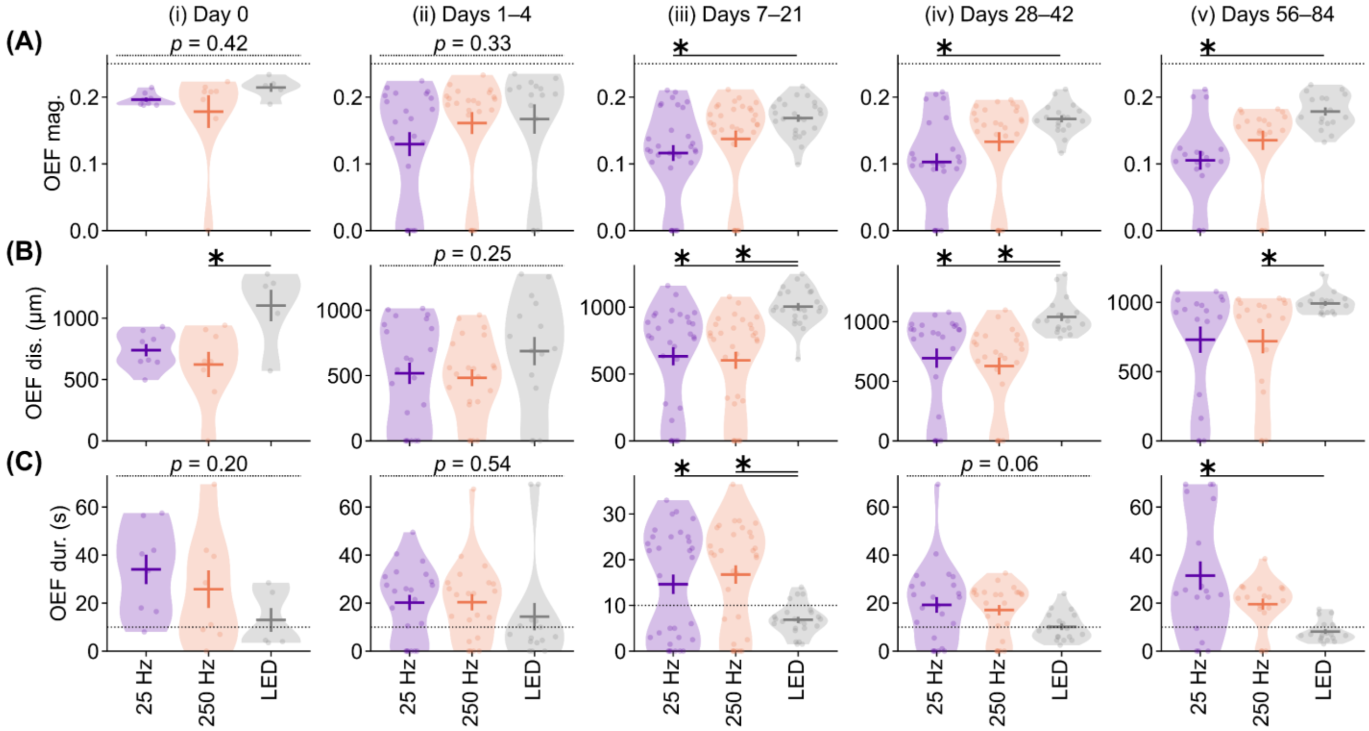


**Figure S8. Temporal grouping reveals sustained recovery of hemodynamic activation and metabolic responsiveness over chronic implantation.** (A–C) Comparisons of OEF activation (A) magnitude, (B) spatial extent (distance), and (C) duration at (i) day 0, (ii) days 1–4, (iii) days 7–21, (iv) days 28–42, and (v) days 56–84 across stimulation conditions. The horizontal dotted line in (A) indicates the pre-stimulation baseline OEF magnitude (0.25), and in (C) indicates stimulation duration (10 s). Horizontal and vertical lines represent mean ± SEM across samples. * with solid line indicates $p < 0.05$ for pairwise comparison using post-hoc Tukey's HSD test. *p* with dotted line indicates *p*-value for one-way ANOVA without significant difference.

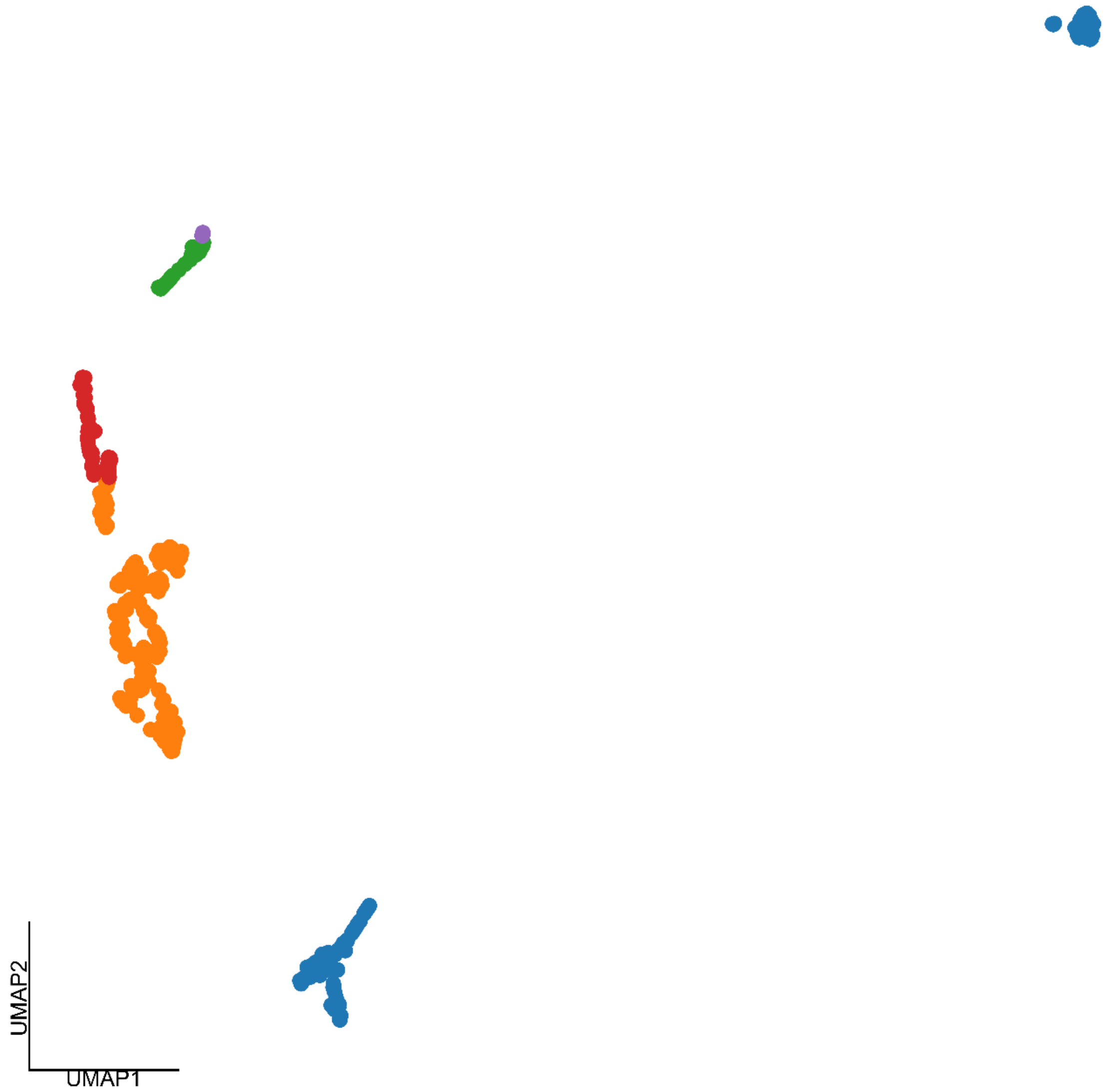


**Figure S9. Dimensionality reduction reveals distinct clustering of epileptiform $Ca^{2+}$ activity from standard activation patterns.** Uniform manifold approximation and projection (UMAP) visualization of $Ca^{2+}$ activation parameters (magnitude, duration, and distance) induced by 25-Hz ICMS. Each symbol represents a single sample from one mouse on one day, projected from three-dimensional feature space into two-dimensional UMAP space using the Python UMAP module. Colors denote clusters identified by agglomerative clustering in the original three-dimensional space (see Fig. 7C), yielding five discrete groups. The epileptiform cluster (green; magnitude > 100%, duration > 20 s, distance > 800 μm) was proximal to, yet clearly separated from, the standard-activation cluster (red), indicating both shared features and distinct network-level dynamics underlying epileptiform events.

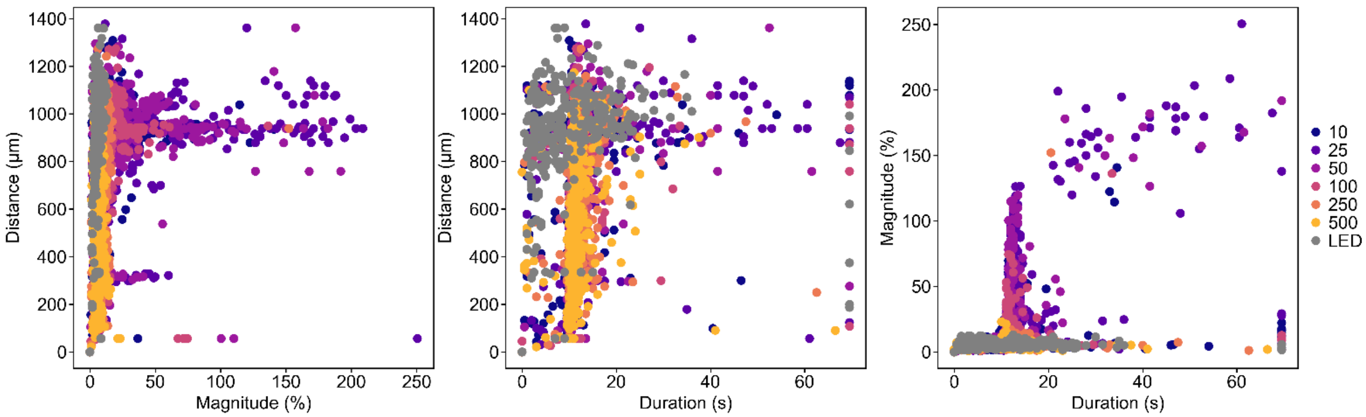


**Figure S10. Epileptiform $Ca^{2+}$ activity occurs predominantly under chronic 25-Hz ICMS stimulation.** Distributions of $Ca^{2+}$ activation magnitude, duration, and distance across all stimulation conditions and time points (0–84 days post-implantation). The epileptiform cluster (magnitude > 100%, duration > 20 s, distance > 800 μm; defined by agglomerative clustering) is composed primarily of data from the 25-Hz ICMS condition, with only a few scattered samples from other ICMS frequencies. These results indicate that chronic 25-Hz ICMS almost exclusively evokes large, long-lasting, and spatially widespread epileptiform activity compared to higher-frequency ICMS or visual stimulation.

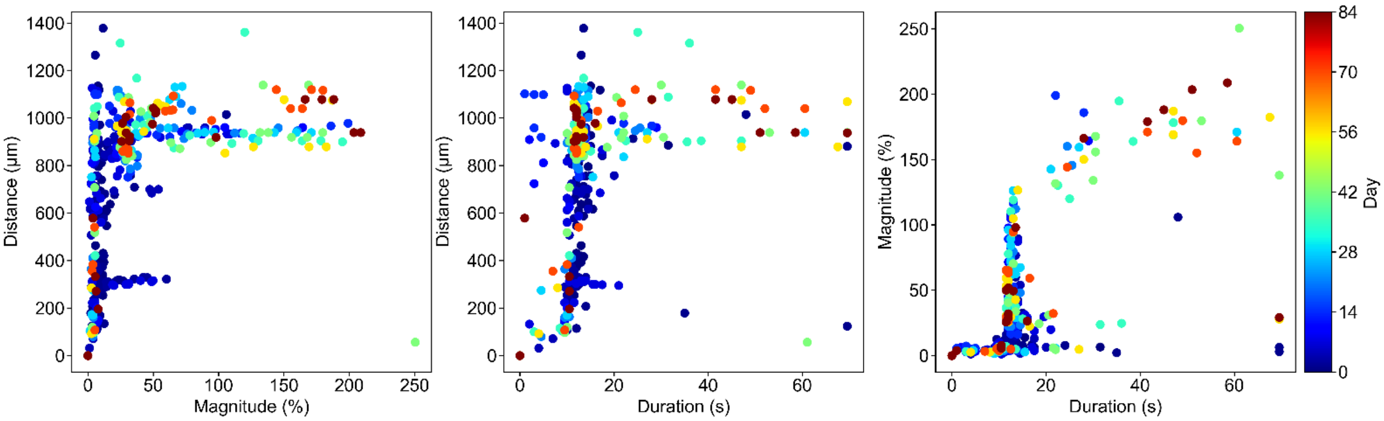


**Figure S11. Epileptiform $Ca^{2+}$ activity emerges predominantly during late chronic days after 25-Hz ICMS.** Distributions of $Ca^{2+}$ activation magnitude, duration, and distance induced by 25-Hz ICMS across post-implantation days. The epileptiform cluster (magnitude > 100%, duration > 20 s, distance > 800 µm; defined by agglomerative clustering) mainly comprises data from late chronic periods (approximately 14–84 days post-implantation, light blue to reddish colors), indicating that epileptiform activity predominantly arises weeks after probe insertion.

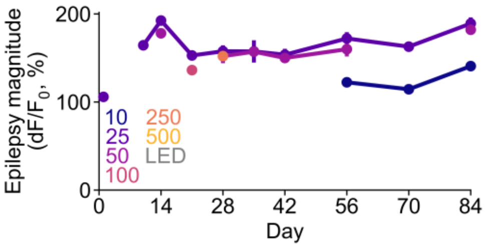


**Figure S12. Magnitude of epileptiform $Ca^{2+}$ activity remains stable across chronic days.** Mean ± SEM of epileptiform activity magnitude over post-implantation days for all stimulation conditions. Only cases exhibiting epileptiform activity (magnitude > 100%, duration > 20 s, distance > 800 μm; defined by agglomerative clustering) are included; data points without epileptiform events are omitted. Magnitudes remained relatively constant across chronic days, indicating that the severity of individual seizure-like events is largely independent of time after surgery and may instead depend on stimulation frequency (e.g., differences between 10 Hz and 25 Hz during days 56–84).

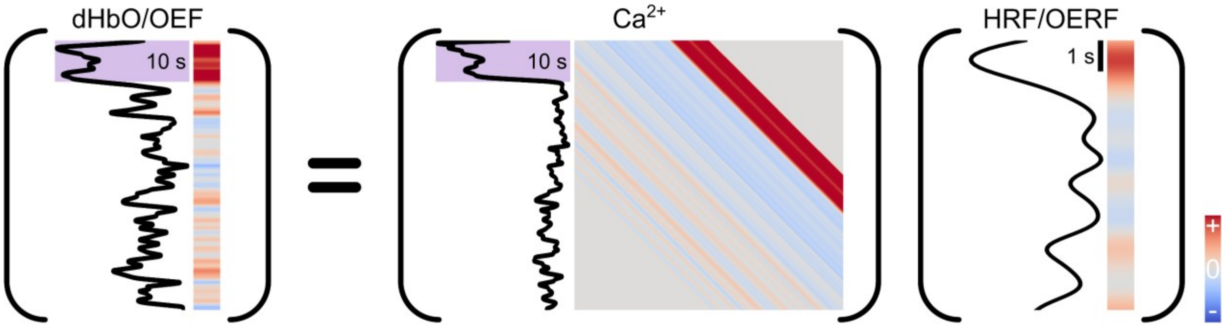


**Figure S13. Derivation of the hemodynamic response function (HRF/OERF) as a measure of neurovascular coupling.** Schematic illustrating the mathematical relationship between neural activity ($Ca^{2+}$), hemodynamic signals (dHbO or OEF), and the conversion function (HRF/OERF). Hemodynamic signals, recorded at the mesoscopic scale (column vector), are modeled as the convolution of neural activity, recorded under mesoscopic-scale imaging (matrix), with the HRF/OERF (unknown column vector). Deconvolution was performed to estimate HRF/OERF, providing a functional measure of neurovascular coupling (see 2.4.5 for details).

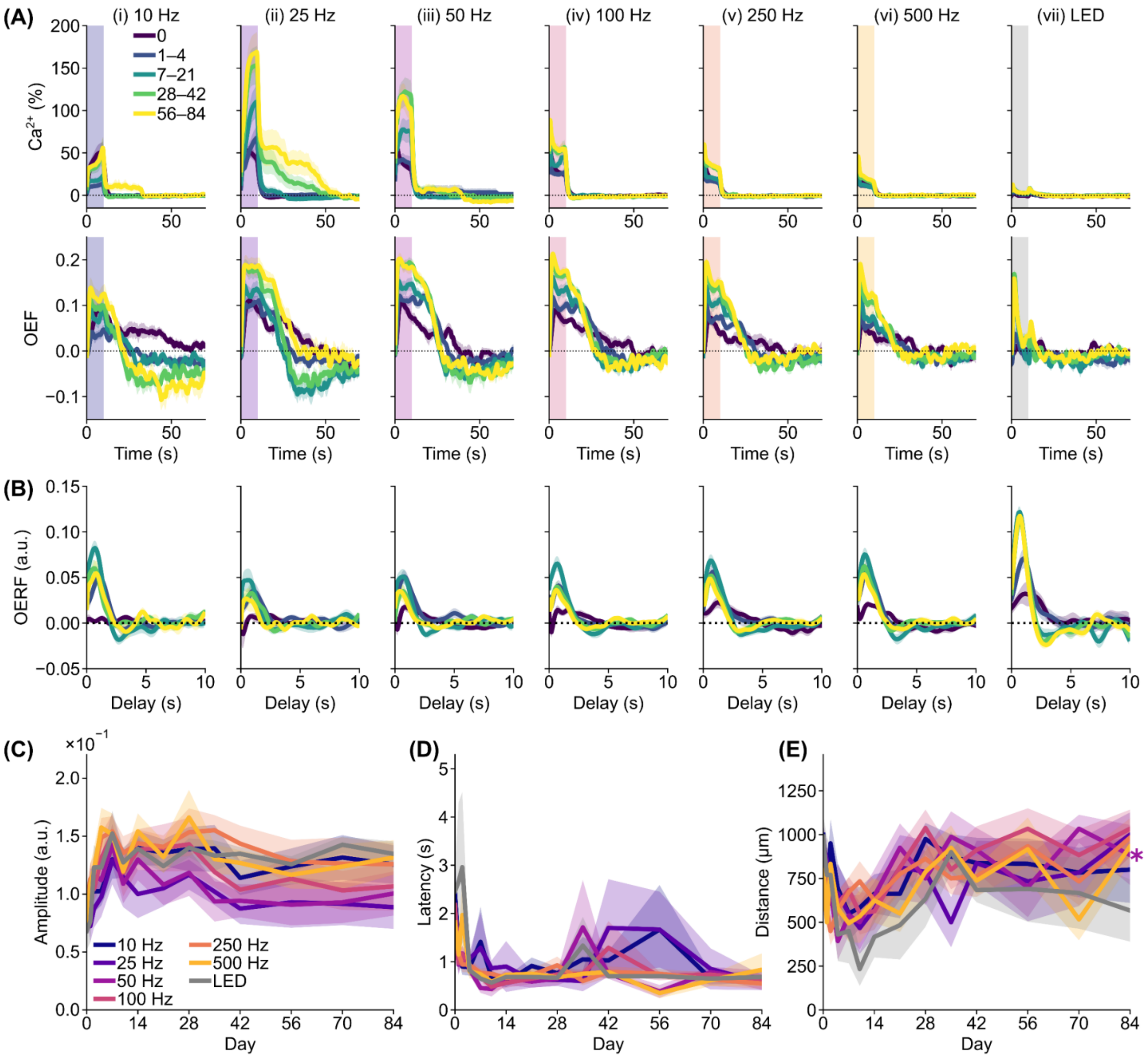


**Figure S14. Progressive recovery of neurovascular coupling following probe insertion.** (A) $Ca^{2+}$ and OEF time courses within 100 μm of the electrode (mean ± SEM across mice). OEF signals were zero-meaned and flipped for visualization. (B) Corresponding OERF profiles derived from deconvolution. (C–E) Comparisons across days post-implantation of (C) peak amplitude, (D) peak latency, and (E) peak distance. Mean ± SEM across mice; * nearby line plot indicates a significant effect of day (LME model).

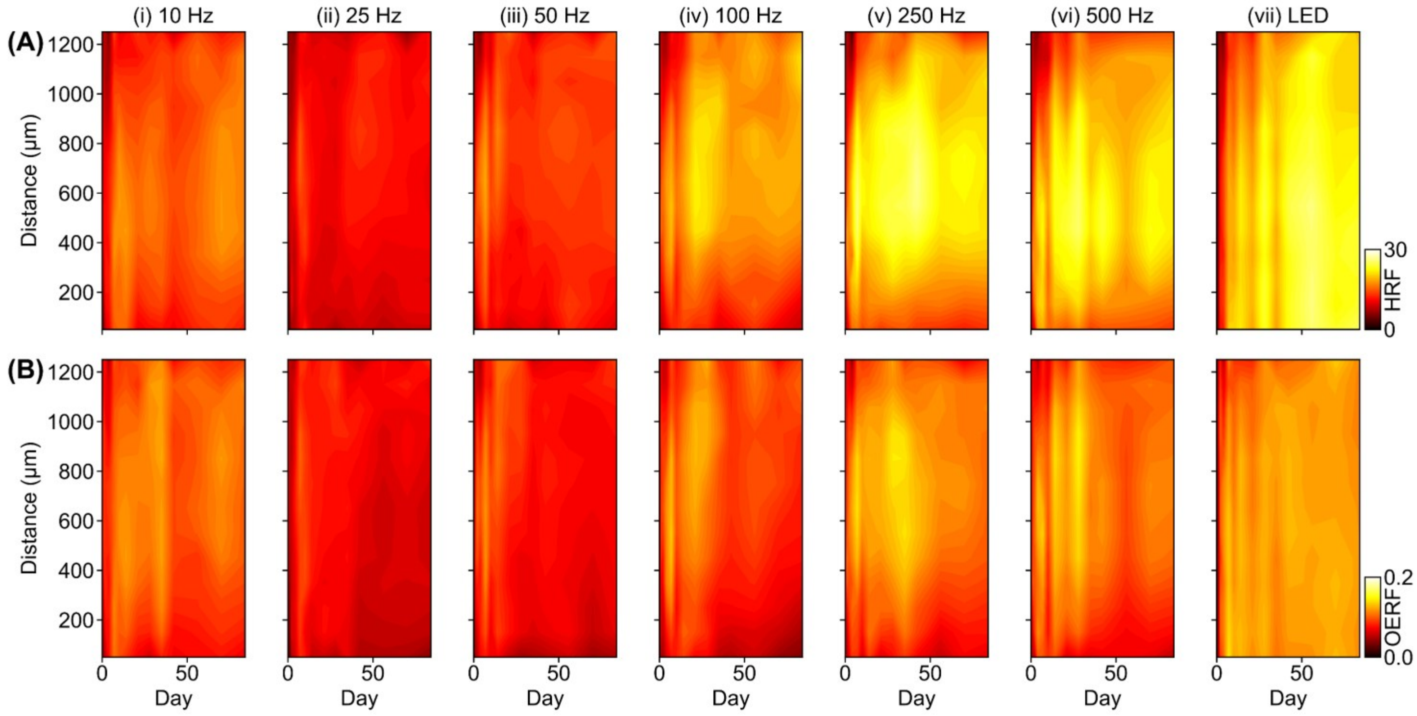


**Figure S15. Spatial and temporal attenuation of hemodynamic response functions near the probe over chronic days.** (A) Peak amplitude of HRF reconstructed from $Ca^{2+}$ and dHbO signals. (B) Peak amplitude of OERF reconstructed from $Ca^{2+}$ and OEF signals. Both metrics show similar trends: reduced peak amplitudes (darker colors) in the immediate vicinity of the probe during ICMS, particularly at chronic time points, whereas local attenuation is less pronounced under visual stimulation.

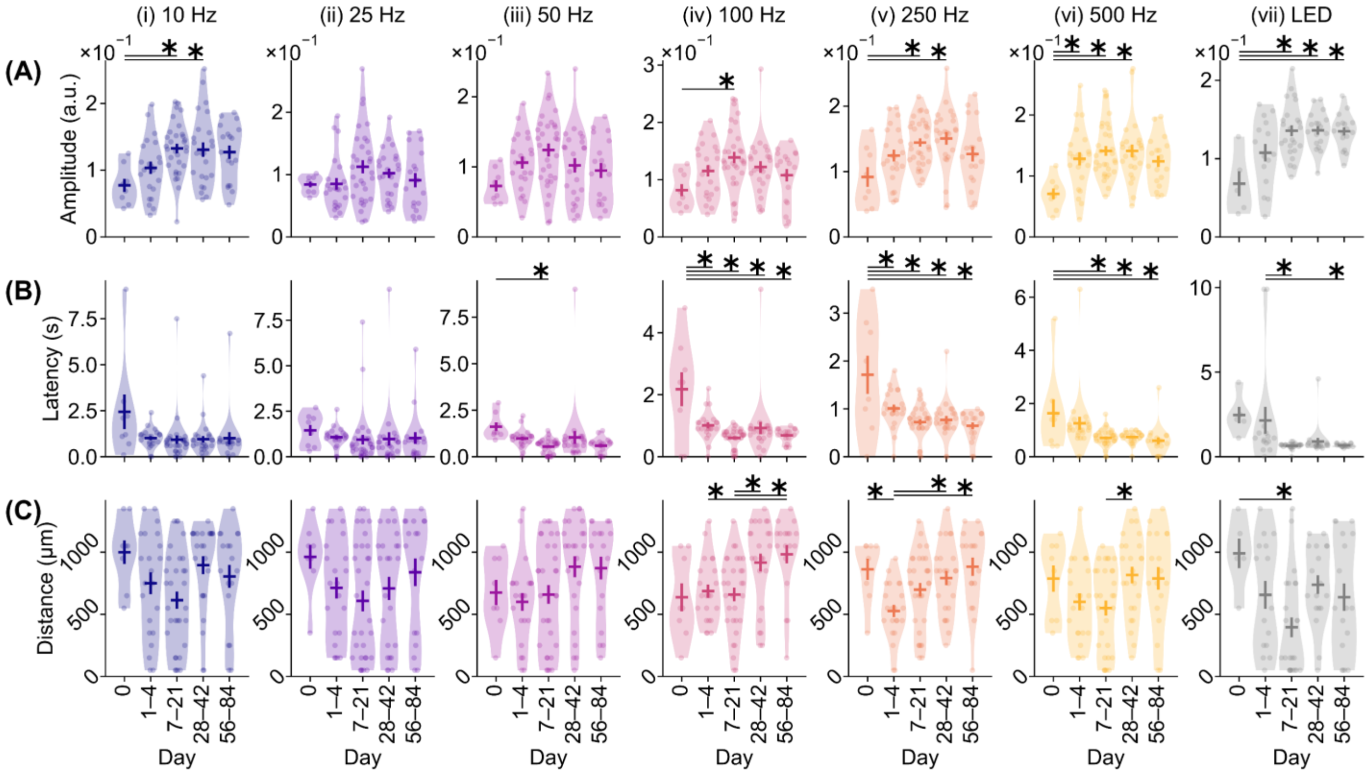


**Figure S16. Chronic evolution of hemodynamic response function characteristics following probe insertion.** Comparisons of OERF (A) peak amplitude, (B) latency to peak, and (C) distance to peak across days after probe insertion. Horizontal and vertical lines indicate mean ± SEM across samples. * with solid line indicates significant differences based on post-hoc Tukey's HSD test.
